# Unraveling the kinetochore nanostructure in *Schizosaccharomyces pombe* using multi-color single-molecule localization microscopy

**DOI:** 10.1101/2021.12.01.469981

**Authors:** David Virant, Ilijana Vojnovic, Jannik Winkelmeier, Marc Endesfelder, Bartosz Turkowyd, David Lando, Ulrike Endesfelder

**Affiliations:** Department of Systems and Synthetic Microbiology, Max Planck Institute for Terrestrial Microbiology and LOEWE Center for Synthetic Microbiology (SYNMIKRO), Marburg, Germany; Department of Physics, Carnegie Mellon University, Pittsburgh, Pennsylvania, US; Institute for Microbiology and Biotechnology, Rheinische-Friedrich-Wilhelms-Universität Bonn, Bonn, Germany; Institute for Assyriology and Hittitology, Ludwig-Maximilians-Universität München, Germany; Department of Biochemistry, University of Cambridge, Cambridge CB2 1EW, UK

**Author notes:** authors contributed equally (David Virant, Ilijana Vojnovic, Jannik Winkelmeier).

## Abstract

The key to ensuring proper chromosome segregation during mitosis is the kinetochore complex. This large and tightly regulated multi-protein complex links the centromeric chromatin to the microtubules attached to the spindle pole body and as such leads the segregation process. Understanding the architecture, function and regulation of this vital complex is therefore essential. However, due to its complexity and dynamics, only its individual subcomplexes could be studied in high-resolution structural detail so far.

In this study we construct a nanometer-precise *in situ* map of the human-like regional kinetochore of *Schizosaccharomyces pombe (S. pombe)* using multi-color single-molecule localization microscopy (SMLM). We measure each kinetochore protein of interest (POI) in conjunction with two reference proteins, cnp1^CENP-A^ at the centromere and sad1 at the spindle pole. This arrangement allows us to determine the cell cycle and in particularly the mitotic plane, and to visualize individual centromere regions separately. From these data, we determine protein distances within the complex using Bayesian inference, establish the stoichiometry of each POI for individual chromosomes and, consequently, build an *in situ* kinetochore model for *S.pombe* with so-far unprecedented precision. Being able to quantify the kinetochore proteins within the full *in situ* kinetochore structure, we provide valuable new insights in the *S.pombe* kinetochore architecture.

## Introduction

The proper segregation of chromosomal DNA during cell division is one of the most crucial processes in the cell cycle of living organisms. Aneuploidy, caused by chromosome maldistribution, leads to cancer, birth defects and cell death (Pfau and Amon, 2012; Santaguida and Amon, 2015; Yuen et al., 2005). During chromosome segregation, sister chromatids are separated by the microtubules of the spindle apparatus. Here, the kinetochore, a multiprotein complex, acts as the link between the centromeric DNA and the microtubules emerging from the spindle (Musacchio and Desai, 2017; Roy et al., 2013).

In budding yeast *Saccharomyces cerevisiae,* the kinetochore appears as a structure of about 126 nm in diameter containing a large central hub surrounded by multiple outer globular domains (Gonen et al., 2012). A single kinetochore connects one single microtubule to a 125 base-pair region, a so-called point centromere of defined sequence (Clarke, 1998; Winey et al., 1995). In contrast, in the fission yeast *Schizosaccharomyces pombe,* the kinetochore complex links 2-4 kinetochore microtubules (kMTs) to a regional centromere in the kilobasepair range (Chikashige et al., 1989; Ding et al., 1993). In higher eukaryotes, the structure is even more extensive. For example, for human chromosomes, the centromeric region spans 0.5-1.5 megabase pairs and is built in α-satellite repeats (Wevrick and Willard, 1989; Zinkowski et al., 1991). Accordingly, the human kinetochore structure allows attachment to approximately 20 kMTs (McEwen et al., 1998; McEwen et al., 1997).

Despite these differences in centromere length and number of kMTs, the general kinetochore architecture is highly conserved (Drinnenberg et al., 2016; van Hooff et al., 2017). The position of the centromeric region is epigenetically defined by a H3 histone variant, cnp1^CENP-A^, which replaces one or both H3 proteins in the centromeric nucleosomes (Dunleavy et al., 2011; Lando et al., 2012). (In this report, the terminology of *S. pombe* is used, with the human homolog in superscript, if different.) cnp1^CENP-A^ provides the scaffolding for the assembly of the kinetochore, which consists of several subcomplexes: the CCAN network, the Mis12/MIND complex, the KNL1 complex, the NDC80 complex and the DAM1/DASH complex (or Ska complex in humans) (Kixmoeller et al., 2020; Musacchio and Desai, 2017).

The structures of these subcomplexes have been made available by cryo-EM and x-ray crystallography studies, e.g. the structures of the CENH3 nucleosome (Migl et al., 2020; Tachiwana et al., 2011), the CCAN network (Hinshaw and Harrison, 2019a; Hinshaw and Harrison, 2019b; Yan et al., 2019), the MIND/Mis12c (Dimitrova et al., 2016; Petrovic et al., 2016), the NDC80c (Ciferri et al., 2008; Valverde et al., 2016) and the Dam1/DASH complex (Jenni and Harrison, 2018). Nevertheless, despite recent advances in reconstituting large sub-complexes, e.g. success in resolving the structure of the 13-subunit Ctf19c/CCAN with a 4 Å resolution *in vitro* (Hinshaw and Harrison, 2019a), the assignment of kinetochore elements to specific subcomplexes within the fully assembled kinetochore structure has not been successful so far (Gonen et al., 2012; Walstein et al., 2021).

Fluorescence microscopy can visualize selected proteins of interest (POIs) using fluorescent labels such as fluorescent proteins (FPs) (Tsien, 1998). Fluorescent labels can be used quantitatively and as such can yield kinetochore protein copy numbers (Coffman et al., 2011; Dhatchinamoorthy et al., 2017; Joglekar et al., 2008; Joglekar et al., 2006; Johnston et al., 2010; Lawrimore et al., 2011; Schittenhelm et al., 2010; Suzuki et al., 2015) or intra-complex distances (Aravamudhan et al., 2014; Haase et al., 2013; Joglekar et al., 2009; Schittenhelm et al., 2007; Suzuki et al., 2014; Suzuki et al., 2018; Wan et al., 2009).

However, the spatial resolution of fluorescence microscopy is not sufficient to resolve individual kinetochores (Tournier et al., 2004). Here, super-resolution microscopy (SRM) bridges the resolution gap between conventional fluorescence microscopy and electron microscopy. The kinetochore has been studied by structured illumination microscopy (SIM) (Dhatchinamoorthy et al., 2019; Schubert et al., 2020; Venkei et al., 2012; Wynne and Funabiki, 2016; Zielinska et al., 2019) and by stimulated emission depletion (STED) microscopy (Drpic et al., 2018; Zielinska et al., 2019). Both SRM techniques provided detailed pictures of the structure at about 100 nm resolution. Only a handful of studies have been carried out at single molecule resolution, so far. Two *in situ* studies each target a single kinetochore POI, cnp1^CENP-A^ (Lando et al., 2012) and Ndc80 (Wynne and Funabiki, 2016), and one study focuses on two-color POI pairs *in vitro* (Ribeiro et al., 2010).

In this study, we investigate the architecture of the regional kinetochore of the fission yeast *S. pombe.* We created a triple-color *S. pombe* strain library of fluorescent fusions of key proteins of the inner and outer kinetochore. Using quantitative multi-color single-molecule localization microscopy (SMLM) imaging and Bayesian inference, we report *in situ* inner-kinetochore distances to the centromere as well as protein copy numbers per centromeric cluster for each POI imaged. With those numbers at hand, we can determine the protein architecture of the kinetochore during mitosis and build an *in situ* kinetochore model at a nanometer resolution. Our study confirms *in vitro* studies on reconstituted kinetochore subcomplexes and adds new *S. pombe* specific insights to our current kinetochore architecture knowledge.

## Materials and methods

### Strain construction

To construct a dual-color reference strain with both a reference at the centromere and at the spindle pole body (SPB), we added a spindle reference to the *S. pombe* strain DL70, which already carries a PAmCherry1-cnp1^CENP-A^ N-terminal fusion (Lando et al., 2012). To tag the SPB protein sad1 with mScarlet-I (Bindels et al., 2017) at its C-terminus, we adapted a cloning strategy from Hayashi et al. (Hayashi et al., 2009): DL70 cells were made chemically competent (Frozen-EZ Yeast Transformation II Kit, Zymo Research, #T2001) and were transformed with a linear DNA fragment consisting of the mScarlet-I sequence and a hygromycin resistance cassette flanked by 500-600 bp of homology arms on both sides for homologous recombination into the genome downstream of the native sad1 gene. Additionally, we inserted the terminator ADH1 from *S. cerevisiae* (Curran et al., 2013) between POI-FP and resistance genes to ensure native protein expression independent from the resistance gene. Cells were streaked on a YES agar plate, incubated overnight at 32°C and replica-plated on a YES agar plate containing 100 mg/L hygromycin B (Roche, #10843555001). The hygromycin plates were incubated for 1-2 days at 32°C until colonies had formed. Colonies were checked for successful integration of the mScarlet-I ADH1 hygromycin cassette by colony PCR and subsequent sequencing. When we tested the resulting strain SP129 for single-molecule imaging, we noted a high level of autofluorescence that we could attribute to an accumulating precursor in the interrupted adenine biosynthesis pathway due to the ade6-210 mutation, which is as bright as the single-molecule FP intensities and therefore severely compromises the SMLM read-out (Supplementary Figure S1 and Supplementary Text S1) (Allshire, 1995; Levenberg and Buchanan, 1957; Lukens and Buchanan, 1959). We thus repaired the ade6 gene using a wildtype strain template. Transformed cells were streaked on EMM 5S agar plates and grown overnight at 32°C, replica-plated on EMM agar plates containing 225 mg/L of histidine, leucine, lysine and uracil each, but only 10 mg/L of adenine and incubated for 1-2 days at 32°C. Cells that still possessed the truncated ade6 gene turned pink due to accumulation of the precursor (which is also used as a selection marker in the literature (Allshire, 1995)). Colonies with a successfully integrated wildtype ade6 gene stayed white and were checked by sequencing. This new strain (SP145) was used as the parent strain for the construction of a three-color kinetochore library: Here, each POI was tagged with the FP mEos3.2-A69T using the same procedure as for sad1-tagging but using a kanamycin resistance cassette (Supplementary Figure S2 a). Successful transformants were selected on YES agar plates containing 100 mg/L geneticin (Roche, #4727894001) and checked with colony PCR and sequencing. Sequencing was done with the Mix2Seq kit (Eurofins Genomics, Germany). A colony was picked for each newly generated strain, cultured to early stationary phase in rich YES medium, mixed in a 1:1 ratio with sterile 87% glycerol and stored at −80°C. All strains used in this study can be found in Supplementary Table S1 and all primers used for strain construction can be found in Supplementary Table S2.

### Spot tests

To assess the effect of the FP tags on the POIs and reference proteins, spot tests were conducted to compare the temperature and thiabendazole (TBZ, inducing mitotic defects by microtubule depolymerization) sensitivity to the parental h^+^ WT strain (Supplementary Figure S3). For this, *S. pombe* strains from cryostocks were streaked on YES agar plates and incubated at 32°C for two days. After growing the overnight cultures in YES medium at 25°C to an OD_600_ of 1-2, all cultures were adjusted to an OD_600_ of 1.0 and 2 μL each of a serial dilution of 1:1, 1:10, 1:100 and 1:1000 were plated either on YES agar plates and incubated at 25, 32 or 37°C for three days for temperature sensitivity testing, or on YES agar plates containing 2.5, 5.0, 7.5 or 10.0 μg/mL TBZ (Cayman Chemical Company, #23391) and incubated at 25°C for three days.

### Flow Cytometry

To assess the effect of the fluorescent protein tags on the phenotype of cells, flow cytometry was conducted (Supplementary Figure S4). Cells were streaked from cryostocks on YES agar plates and grown at 32°C for 2-3 days until colonies were visible. A single colony was picked, inoculated in 10 mL rich YES medium and grown overnight (25°C, 180 rpm). Subsequently, the OD_600_ for each strain was measured and an appropriate cell amount was used to inoculate 10 mL of EMM 5S to the same starting OD_600_. Cells were grown overnight (25°C, 180 rpm) and the OD_600_ was measured again the next morning to verify that cells were in logarithmic growth phase. Light scattering (FSC and SSC) was measured with a BD LSRFortessa flow cytometer (BD Biosciences, Germany). Before each measurement, samples were mixed thoroughly to disperse cell aggregates. For each strain, 10.000 cells were recorded using the FACSDiva software (BD Biosciences, Germany). Data was analyzed and plotted with the software FlowJo (BD Biosciences, Germany).

### Sample preparation for SMLM imaging

After streaking *S. pombe* strains from cryostocks on YES agar plates and incubating them at 32°C until colonies were visible, overnight cultures were inoculated in rich YES media at 25°C and grown to an OD_600_ of 1-2. Next, 2x 50 mL night cultures were inoculated in freshly prepared EMM 5S media starting at OD_600_ 0.1 and grown at 25°C for around 16 h to OD_600_ 1.0-1.5. Cultures were centrifuged at 2000 rpm at 25°C for 5 min, the supernatant was discarded, and the pellets were resuspended in 200 μL fresh EMM 5S each. Meanwhile, for cell synchronization, two previously prepared and frozen 15 mL falcons containing 20% lactose (Carl Roth #6868.1) were thawed at 30°C and the cell suspension was carefully added on top of the still cold lactose gradient (Supplementary Figure S2 b). The falcons were then centrifuged at 1000 rpm for 8 min and 1 mL of the resulting upper cell layer, which consists of cells in early G2 stage, was extracted, added to 1 mL fresh EMM 5S and centrifuged at 4000 x g for 1 min. The pellets were resuspended in 500 μL EMM 5S, combined and the OD_600_ was measured. The cell suspension was inoculated in fresh EMM 5S at a starting OD_600_ of 0.6 and grown for 60, 70 or 80 min at 25°C. At each time point, 2 mL aliquots were taken from the main culture for chemical fixation with 3% paraformaldehyde (PFA, Sigma-Aldrich, #F8775) at 20 min at RT. The fixed cells were centrifuged at 4000xg for 2 min and washed six times with 1 mL washing buffer alternating between 1xPBS, pH 7.5 (Gibco, #70011-036) and 50 mM Tris-HCl, pH 8.5 (Carl Roth, #AE15.2) with each 10 min of incubation. The cells were resuspended in 500 μL Tris-HCl after the last wash, a 1:1000 dilution of previously sonicated dark red (660/680) FluoSpheres (Molecular probes, #F8807) was added to serve as fiducial markers for drift correction (Balinovic et al., 2019) and incubated for 10 min in the dark. Ibidi 8-well glass bottom slides (Ibidi, #80827) were cleaned by incubating 1M KOH for 30 min, coated with poly-L-lysine (Sigma-Aldrich, #P8920-100ML) for 20 min and air dried thereafter. Next, 200 μL of the bead/cell mix were added to each well and were incubated for 10 min in the dark. Meanwhile, a 0.5% w/v low gelling agarose solution (Sigma-Aldrich, #A9414-100G) was melted in 50 mM Tris-HCl at 70°C for 1-2 min, a 1:1000 dilution of dark red (660/680) FluoSpheres was added, and the mix was kept at 32°C until further use. The supernatant of the bead/cell mix incubating in the Ibidi 8-well glass bottom slide was carefully removed on ice and six drops of the agarose mix were carefully added on each well and incubated for 5 min for proper gelation. Finally, 200 μL 50 mM Tris-HCl, pH 8.5 were added to each well. Samples were stored at 4°C for a maximum of one day until being imaged.

### Microscope setup and SMLM Imaging

A custom built setup based on an automated Nikon Ti Eclipse microscopy body including suitable dichroic and emission filters for SMLM imaging using Photoactivated Localization Microscopy imaging by primed photoconversion (PC-PALM) (ET dapi/Fitc/cy3 dichroic, ZT405/488/561rpc rejection filter, ET610/75 bandpass, all AHF Analysentechnik, Germany) and a CFI Apo TIRF 100×oil objective (NA 1.49, Nikon) was used for multi-color imaging experiments as described earlier (Virant et al., 2017). The Perfect Focus System (PFS, Nikon, Germany) was utilized for z-axis control and all lasers (405 nm OBIS, 488 nm Sapphire LP, 561 nm OBIS, Coherent Inc. USA) except the 730 nm laser (OBIS, Coherent Inc. USA) were modulated by an acousto-optical tunable filter (AOTF) (TF525-250-6-3-GH18, Gooch and Housego, USA) in combination with the ESio AOTF controller (ESImaging, UK). Singlemolecule fluorescence was detected using an emCCD camera (iXON Ultra 888, Andor, UK) at a pixel size of 129 nm and the image acquisition was controlled by a customized version of μManager (Edelstein et al., 2010).

Movie acquisition of all triple-color strains was performed sequentially in HILO mode (Tokunaga et al., 2008). First, the sample was illuminated at low intensity 561 nm with 0.1-1 W*cm^-2^ at the sample level to identify fields of view containing a good number of cells with two SPB spots (metaphase and early anaphase A cells) aligned in parallel to the focal plane. The sad1-mScarlet-I signal was read-out by taking 100 images of 60 ms exposure time with the 561 nm laser at 200 W*cm^-2^ (at the sample level). Residual mScarlet-I signal was photobleached using 1 kW*cm^-2^ of 561 nm laser light for 30-60 s. Next, the green-to-red photoconvertible FP mEos3.2-A69T was imaged in the same spectral channel using primed photoconversion by illuminating the sample with 488 (at every 10th frame) and continuous 730 nm laser light (6 - 70 & 450 W*cm^-2^ respectively) and read-out by 561 nm at 1 kW*cm^-2^ for 15.000 frames of 60 ms exposure. An additional post-bleaching step with 488 nm laser light (210 W*cm^-2^ for 20 s) was performed to ensure that no residual mEos3.2-A69T signal was detected in the next acquisition step. Finally, PAmCherry1-cnp1^CENP-A^ was photoactivated by illumination with 10 - 20 W*cm^-2^ of 405 nm laser light and read-out by 561 nm at 1 kW*cm^-2^ for 10.000 frames of 60 ms exposure. Keeping the field of view well defined, we ensured homogeneous illumination.

### Staining of samples with organic dyes

The construction of the N-terminal Halo-cnp1 strain is described in (Vojnovic, 2016). An overnight Halo-cnp1 YES culture was harvested by centrifugation, resuspended in fresh YES and fixed with 3.7% PFA at room temperature (RT) for 10 min. Afterwards, the sample was washed 3x and residual PFA was quenched using 1x PEM (100 mM PIPES (Sigma-Aldrich, #P1851-100G), pH 6.9, 1 mM EGTA (Sigma-Aldrich, #EDS-100G), 1 mM MgSO4 (Sigma-Aldrich, #M2643-500G)) containing 50 mg/mL NH4Cl (Carl Roth, #P726.1). Next, the sample was permeabilized using 1.25 mg/mL Zymolyase (MP Biomedicals, #320921) in 1x PEM at 37°C for 10 min and subsequently washed 3x with 1x PEM for 10 min. The sample was incubated using a few drops of Image-iT FX signal enhancer (Invitrogen, #I36933) at RT for 1 h to prevent non-specific staining and afterwards stained with 50 nM Halo-CF647 in PEMBAL buffer (1x PEM containing 3% BSA (Sigma-Aldrich, #A8549-10MG), 0.1% NaN_3_ (Carl Roth, #4221.1), 100 mM lysine hydrochloride (Sigma-Aldrich, #L5626)) at RT for 3 h. The stained cells were then washed four times for 10 min with alternating 1x PEM and 1x PBS and incubated at RT for 15 min on a previously washed and with poly-L-lysine coated Ibidi 8-well glass bottom slide. Finally, the attached cells were washed twice with 1x PEM for 10 min.

### SMLM imaging and post-processing of dye-stained samples

The stained Halo-cnp1 strains were imaged in a redox buffer (100 mM MEA (Sigma-Aldrich, #MG600-25G) with an oxygen scavenger system (van de Linde et al., 2011) in 1x PEM) and illuminated with a 637 nm laser (OBIS, Coherent Inc. USA) at 1.4 kW*cm^-2^ at the sample level with an exposure time of 50 ms for 10.000 frames on the same setup as described before, except that a different dichroic (ZET 405/488/561/640, Chroma) and longpass filter (655 LP, AHF) were used. Single-molecule localizations were obtained by the open-source software Rapidstorm 3.2 (Wolter et al., 2012). In order to track localizations that appeared in several frames and thus belong to the same emitting molecule, the NeNA localization precision value (Endesfelder et al., 2014) was calculated in the open-source software Lama (Malkusch and Heilemann, 2016).

### Reconstruction of SMLM images

For visualization, we aimed to reconstruct SMLM images that neither over- nor under-interpret the resolution of the SMLM data and resemble fluorescence images as closely as possible. Localizations were tracked together using the Kalman tracking filter in Rapidstorm 3.2 with two sigma, and the NeNA value used as sigma (dSTORM data), or using kd-tree tracking (3Dcolor strain library data, described in detail in the data analysis section below). Images were then reconstructed in Rapidstorm 3.2 or 3.3 with a pixel size of 10 nm. Rapidstorm linearly interpolates the localizations and fills neighboring pixels based on the distance between the localization and the center of the main subpixel bin to avoid discretization errors (Wolter et al., 2012). These images were then processed with a Gaussian blur filter based on their NeNA localization uncertainty in the open-source software ImageJ 1.52p (Schindelin et al., 2012). Importantly, images were only used for image representation purposes, all data analysis steps were conducted on the localization data directly (see data analysis).

### Construction of the FtnA protein standard

The pRSETa-FtnA backbone was amplified from pRSETa BC2-Ypet-FtnA (Virant et al., 2018), the mEos3.2-A69T fragment was amplified from pRSETa-mEos3.2-A69T (Turkowyd et al., 2017) with a 18 and 23 bp overlap to the pRSETa-FtnA backbone respectively. Purified DNA fragments (Clean & Concentrator-5 Kit, Zymo Research, #D4013) were merged by Overlap-Extension PCR in a 1:2 ratio of backbone to insert. The resulting plasmid was purified and transformed into competent DH5α cells and incubated on LB AmpR agar plates at 37°C overnight. Plasmids were isolated from individual colonies (Monarch Plasmid Miniprep Kit, New England Biolabs, #T1010L) and sequenced (Mix2Seq Kit NightXpress, Eurofins genomics). The confirmed construct was transformed into competent Rosetta DE3 cells.

### Sample preparation and imaging of the FtnA protein standard

Rosetta DE3 containing pRSETa mEos3.2-A69T-FtnA were grown over night at 37°C and 200 rpm in 50 mL AIM (autoinducing medium containing freshly added 25 g/L glucose (Carl Roth, #X997.2) and 100 g/L lactose (Carl Roth, #6868.1)). Cells were harvested by centrifugation at 4000 x g and 4°C for 30 min and the pellet was resuspended in 2 mL 50 mM Tris HCl (Carl Roth, #AE15.2) containing 25 mM NaCl (Sigma-Aldrich, #S3014-500G) at pH 8.5. The bacterial cell wall was digested by incubation with 50 mg/mL Lysozyme (Sigma-Aldrich, #L2879) for 2 h at 4°C. The sample was homogenized (Tough micro-organism lysing tubes (Bertin, #VK05) with 0.5 mm glass beads in Precellys 24 tissue homogenizer at 3x 15 s and 5000 rpm at RT). The cell lysate was centrifuged at 4000 x g for 30 min at 4°C and the supernatant was transferred into a clean Eppendorf tube. The centrifugation and transfer were repeated to ensure a debris-free supernatant. For long-term storage 30% glycerol (Sigma-Aldrich, #G5516) was added and the stock was kept at −20°C until further use.

In order to prepare the FP-FtnA oligomer surfaces, the mEos3.2-A69T-FtnA stock was diluted 1:5000 in 50 mM Tris HCl containing 25 mM NaCl at pH 8.5 and incubated with a sonicated 1:1000 dilution of dark red (660/680) FluoSpheres (Molecular probes, #F8807) on a clean, poly-L-lysine coated Ibidi 8-well glass bottom slide for 10 min. The sample was washed twice using 50 mM Tris HCl containing 25 mM NaCl at pH 8.5 and 0.5% w/v low gelling agarose solution (Sigma-Aldrich, #A9414-100G) was carefully added to the top to ensure proper immobilization. After gelation for 5 min on ice 300 μL of 50 mM Tris HCl containing 25 mM NaCl at pH 8.5 were added to the imaging well.

The same microscope setup and imaging routine that was used for imaging the triple-color strain library was utilized to image the isolated mEos3.2-A69T-FtnA oligomers. To simulate the mScarlet-I imaging & bleaching step prior to mEos3.2-A69T imaging, the sample was first illuminated with the 561 nm laser at 1 kW*cm^-2^ for 1 min before the PC-PALM read-out of mEos3.2-A69T was recorded.

### Data analysis

An overview of the workflow can be found in Supplementary Figure S5.

#### Localization, drift correction and filtering

First, all movies were localized using ThunderSTORM (Ovesny et al., 2014) with the B-spline wavelet filter with a scale of 2.0 pixel and order of 3, a 8 pixel neighborhood and threshold of 1.5*std(wavelet), the integrated Gaussian PSF model with a sigma of 1.4 pixel and a weighted least squares optimization with a fit radius of 4 pixel. The fluorescent beads embedded in the sample were used for drift correction. The combined drift trace as well as the drift trace for each bead were saved and checked for inconsistencies. In those instances, the localization file was manually drift corrected using a custom script written in Python 3 that allowed the selection of individual beads.

A nearest neighbor tracking algorithm based on kd-tree (Jones et al., 2001) was run on the localization files. For each localization, the nearest neighboring localization in a 150 nm radius was identified in the first subsequent frame. Neighbor identifiers were stored and used to connect localizations into tracks. Tracks were then merged by averaging the coordinates, intensity and PSF width of all localizations in a track. Averaged localizations were filtered based on these parameters to discard statistical outliers (Q1 - 1.5*IQR, Q3 + 1.5*IQR) in intensity and chi-square goodness of fit. For the PSF width, a set threshold of a sigma of 70-200 nm was used.

#### Visualization and Manual Analytics

For several manual steps, localizations were visualized in a custom software able to flexibly zoom in/out and to switch between/overlay sad1/POI/ cnp1^CENP-A^ channels. Using this tool, individual localizations could be selected and classified. For channel alignment, localizations belonging to the same fiducial marker in all three channels were grouped together. Cells with visible kinetochore protein clusters in the focal plane were selected and classified as individual region of interests (ROIs) and all clusters were annotated. cnp1^CENP-A^ clusters were paired together with corresponding POI clusters. Whenever there was any doubt whether two clusters belonged to the same kinetochore or whether a cluster represented a single centromere region or several, the clusters were discarded. Two exemplary data sets can be found in the zip-file Supplementary data 1. The annotation work was quality-checked by cross-checking the annotation of two different persons.

#### Channel alignment

The alignment between two channels along a given axis was calculated as the inversevariance weighted mean displacement of beads present in both channels. The position of a bead in each channel was calculated as the mean position of all localizations associated with the given bead in this channel. The variance of the displacement is then given by the sum of the squared standard errors of the two positions. The alignment of two channels was deemed viable if it was based on at least two beads and a minimum of 15000 (channels 1 & 2) or 180 (channel 3) localizations. Initially, only beads with 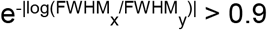 were used.

For all movies, where this did not result in a viable alignment, this threshold was successively lowered to 0.8, 0.7, 0.6, and 0.5 until a viable alignment was found. Movies for which no viable assignment could be found using this procedure were not taken into consideration when estimating the cnp1^CENP-A^-POI distances.

#### Distance calculation

We used Bayesian inference to determine the real distance between cnp1^CENP-A^ and the POI from the measured centers of their respective clusters, assuming Gaussian measurement errors of uncertain size. To improve accuracy, we also took the position of the associated spindle pole into account, as the three points (centers of POI, cnp1^CENP-A^ and corresponding sad1 cluster) can be assumed to lie on a straight line. As it is not possible to reliably determine which spindle pole a centromere is attached to, we built a mixture model to take both possibilities into account. (The SPB closest to a kinetochore is not necessarily the SPB to which the kinetochore is attached to. The reasons for this are that *S. pombe* undergoes closed mitosis where the nuclear envelope doesn’t break up before chromosomes separate and, furthermore, kinetochores oscillate back and forth along the mitotic spindle until DNA segregation takes place (Rieder et al., 1986)). To formalize the idea that a centromere is more likely to be attached to the closer spindle pole, we calculated a mixture coefficient λ = 1 / (1 + (d_2_/d_1_)^-3^), where d_1_ and d_2_ are the measured distances between cnp1^CENP-A^ and the first and second spindle pole, respectively. The coefficient then corresponds to the prior probability that the centromere is attached to the first spindle pole. We visualize the model in Supplementary Figure S6.

We considered three types of Gaussian measurement errors: the measurement error of the spindle, the measurement error of the center of the POI and cnp1^CENP-A^ clusters, and the error of the alignment of the different channels. As the localizations belonging to a single spindle pole are (approximately) normally distributed, the variance of their mean can be estimated with high precision from the sample variance in *x* and *y.* The variances of the errors of the clusters and the alignment are more difficult to estimate directly and were therefore estimated together with the cnp1^CENP-A^-POI distances, using a scaled inverse χ^2^ distribution as prior. Whereas the measurement error can be different for each individual cluster, the alignment error was assumed to be identical for all clusters in a movie. For the alignment error, we estimated the scale parameter *τ*^2^ of the prior distribution from the position of the beads in different channels (although this is likely to be an underestimation, as the same beads were used to calculate the alignment) and the degrees of freedom *v* from the number of beads used in the movie. We used constant values *τ*^2^ = 100 nm^2^ and *v* = 4 for the error of the cluster centers, as they seemed to work reasonably well upon visual inspection. A more objective estimation of this error is difficult, as it results mainly from errors in the clustering, which was performed manually.

We used the Hamiltonian Monte Carlo algorithm as implemented by Stan (Carpenter et al., 2017; Stan Development Team, 2021) and the CmdStanPy python package (Stan Development Team, 2019) to approximate the posterior distribution of the cnp1^CENP-A^-POI distances according to our model (code in Supplementary File 1 and on GitHub: https://github.com/Endesfelder-Lab/Kinetochore_Distances). Posterior means are plotted in Figure 2 and summarized in Table 1, full posterior distributions are shown in Supplementary Figure S7.

**Table 1:** Statistics of distance calculations. For each POI, the posterior mean distance, posterior standard deviation (STD) and the number of kinetochores analyzed is listed. Full posterior distributions, approximated using Hamiltonian Monte Carlo (see Materials & Methods), can be found in Supplementary Figure S7.

| **POI** | **mean distance [nm]** | **STD [nm]** | **N** |
| --- | --- | --- | --- |
| **cnp20**<sup>CENP-T</sup> | 8.3 | 2.4 | 49 |
| **fta2**<sup>CENP-P</sup> | 11.4 | 1.9 | 82 |
| **fta7**<sup>CENP-Q</sup> | 11.7 | 2.9 | 58 |
| **spc7**<sup>KNL1</sup> | 25.9 | 1.2 | 215 |
| **nnf1**<sup>PMF1</sup> | 27.1 | 1.4 | 161 |
| **mis12** | 28.0 | 1.6 | 102 |
| **spc25** | 21.9 | 2.2 | 51 |
| **ndc80**<sup>HEC1</sup> | 36.2 | 1.5 | 135 |
| **dam1** | 64.2 | 1.3 | 155 |

#### Protein Stoichiometry

SMLM localization counts do not directly reflect on the real POI copy numbers due to several over- and undercounting factors (e.g. incomplete FP maturation or photoconversion or FP blinking (Turkowyd et al., 2016)). A robust way to acquire statistically correct average POI copy numbers is to calibrate the localization count data with corresponding data of a protein standard of known stoichiometry using the same sample, imaging and analysis protocols as for imaging the POI. We utilized the *Escherichia coli* protein ferritin (FtnA), a homo-oligomeric protein standard of 24 subunits labeled by mEos3.2-A69T to calibrate our POI localization counts (Finan et al., 2015; Virant et al., 2018). FtnA assembles from monomers into dimers to form 8mers which then arrange into the final 24mer structure. As depicted in Supplementary Figure S8a, FtnA assembly intermediates (FtnA monomers, dimers, 8mers), full 24mers as well as aggregates can be identified by eye and their intrinsic stoichiometric differences are sufficient to discriminate them by fluorescence intensities. We manually selected 1458 8mers and 725 24mers from 29 imaging ROIs from two imaging days. Their localization count distribution yields a mean of 7.27 ± 2.72 STD and 21.68 ± 10.2 STD counts per 8mer and 24mer cluster, respectively (Supplementary Figure S8 b and c). Thus, both give a correction factor of about 0.9 to translate localization counts into POI counts. With the help of this factor, all POI counts were translated into POI copy numbers per centromeric cluster as given in Table 2.

**Table 2:** Statistics of POI localization counts per centromeric region. Distributions can be found in Supplementary Figure S9. *Mean POI copy numbers were derived from the mean localization counts of each POI using 0.9 as a calibration factor as determined by the FtnA protein standard (see Materials & Methods). N number of centromeric regions analyzed.

| POI | mean count | STD | SE of mean | median | Mean POI copy numbers* (± SE) | FP tag | N |
| --- | --- | --- | --- | --- | --- | --- | --- |
| **cnp1**<sup>CENP-A</sup> | 46.8 | 28.0 | 0.8 | 40.0 | ND | PamCherry1 | 1245 |
| **cnp20**<sup>CENP-T</sup> | 55.0 | 32.5 | 4.3 | 47.0 | 61.1 ± 4.8 | mEos3.2-A69T | 57 |
| **fta2**<sup>CENP-P</sup> | 49.6 | 27.9 | 3.0 | 43.0 | 55.1 ± 3.3 | mEos3.2-A69T | 87 |
| **fta7**<sup>CENP-Q</sup> | 51.7 | 28.9 | 3.7 | 46.0 | 57.4 ± 4.1 | mEos3.2-A69T | 61 |
| **spc7**<sup>KNL1</sup> | 85.9 | 48.4 | 3.0 | 76.0 | 95.4 ± 3.3 | mEos3.2-A69T | 259 |
| **nnf1**<sup>PMF1</sup> | 83.3 | 47.2 | 3.2 | 70.0 | 92.6 ± 3.6 | mEos3.2-A69T | 217 |
| **mis12** | 65.8 | 33.1 | 2.8 | 57.5 | 73.1 ± 3.1 | mEos3.2-A69T | 136 |
| **spc25** | 107.2 | 52.2 | 5.6 | 49.5 | 119.1 ± 6.2 | mEos3.2-A69T | 86 |
| **ndc80**<sup>HEC1</sup> | 96.5 | 51.2 | 3.9 | 88.0 | 107.2 ± 4.3 | mEos3.2-A69T | 172 |
| **dam1** | 57.3 | 33.1 | 2.2 | 49.5 | 63.7 ± 2.4 | mEos3.2-A69T | 232 |

### Verifying the reliability of measured localizations counts and distances

In all measurements, we quantitatively assessed not only the localization counts for the POI clusters, but also the counts of the corresponding PAmCherry1-cnp1^CENP-A^ cluster. We use these reference counts to identify possible inconsistencies in the sample preparations and imaging routines of the different experiments. The PAmCherry1 localization counts for the cnp1^CENP-A^ reference remained constant independent of individual sample preparations, imaging days and different strains as can be seen in Supplementary Figure S9 b, where the data is exemplarily sorted into the different strain categories. The POI protein copy numbers as given in Table 2 show a large coefficient of variation. To assess to which extend this variability reflects a technical inability to measure protein levels accurately or some flexibility in kinetochore protein stoichiometry (e.g. due to differing numbers of kMTs per kinetochore), we can use the data of the FtnA oligomer counting standard: The FtnA oligomer is a biologically highly defined structure. Thus, our FtnA measurements can directly serve as a proxy for the contribution of the technical inaccuracy of our PALM imaging and analysis strategy to the variance. Using the results of 21.68 counts ± 10.2 STD for the 24mers and of 7.27 counts ± 2.72 STD for the 8mers, we can estimate that the technical inaccuracy causes a coefficient of variance of 0.35 to 0.5, thus almost completely explaining the experimentally seen coefficient of ~ 0.5 for our POI data (Table 2). Due this high technical inaccuracy, we cannot resolve sub-populations of possibly different kinetochore structures (and thus POI copy numbers) on 2-4 kMTs in our current counting data (Supplementary Figure S9).

For distances, we examined whether the POI-cnp1^CENP-A^ distance depends on the mitotic spindle length, e.g. due to different phases in chromosome separation or different forces. As seen in Supplementary Figure S10 for the POI dam 1 (but being true for all our measurements (data not shown)), we do not detect any dependence of the centroid distance to the mitotic spindle length and therefore conclude that the kinetochores, as we analyze them, all show the same layers and stretching. This is further validated by the posterior distributions of POI- cnp1^CENP-A^ distances in Supplementary Figure S7, which consistently show only a single, well defined peak. Furthermore, using a nup132-GFP strain in early G2 phase, we measured the average nuclear diameter of *S. pombe* to be 2.4 μm ± 0.19 μm (data not shown), in agreement with the literature (Maclean, 1964; Toda et al., 1981). The spindles included in our analysis have spindle lengths well below the nuclear diameter, thus excluding anaphase B cells (Supplementary Figure S10).

## Results & discussion

### A strategy to unravel the kinetochore structure

To investigate the kinetochore architecture during mitosis, we first designed a robust and quantitative multi-color SMLM imaging strategy which did not exist for *S. pombe* before (Vojnovic et al., 2019). Specifically, our aim was to measure protein stoichiometries as well as distances between different POIs in the kinetochore to build a molecular kinetochore map (Figure 1a and b). We planned a triple-color strain library in which each strain contained the same two labeled reference proteins and a varying POI: A spindle reference to identify cells in metaphase/early anaphase and the orientation of the mitotic spindle and a second, centromeric reference to serve as the landmark for kinetochore assembly. To be a reliable reference, the chosen proteins needed a defined organization, sufficient abundance and to be present throughout the cell cycle. This led us to choose sad1, a protein from the SPB (characterized in (Vojnovic, 2016)) and the centromere-specific histone protein cnp1^CENP-A^ (characterized in (Lando et al., 2012)).

**Figure 1:**
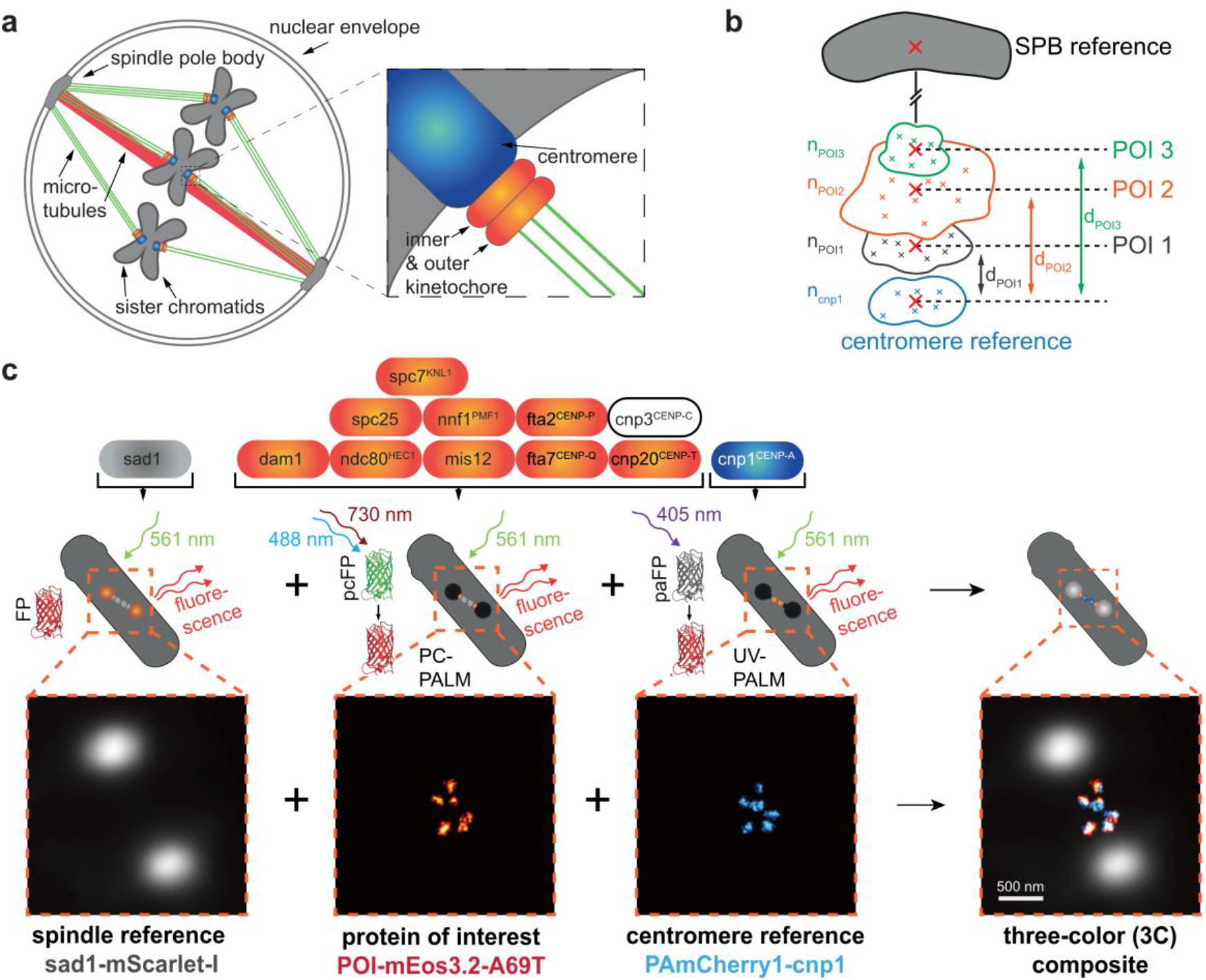
Imaging strategy for measuring the kinetochore nanostructure. **a** During mitosis, the three sister chromatid pairs in *S. pombe* are attached to the two spindle pole bodies (SPBs) by tethering microtubules (green), and the SPBs are pushed apart by a bundle of spindle microtubules (red). The attachment between the centromeric region of each chromatid (blue) and the microtubules is facilitated by the kinetochore-protein complex (orange), which consists of an inner and outer part. **b** Each protein of interest (POI) in the kinetochore complex can be mapped into the kinetochore nanostructure by localizing it in relation to two references, a centromere and an SPB reference. Information about its copy number n_POI_ and its orientation and distance d_POI_ to the reference proteins is obtained. **c** Imaging strategy: First, the SPB protein sad1-mScarlet-I fusion is imaged by conventional epifluorescence microscopy to map the mitotic spindle. Then, both the POI and the centromere reference cnp1^CENP-A^ are measured by super-resolution microscopy: Here, each POI-mEos3.2-A69T fusion (here spc7) is imaged by primed photoconversion PALM (PC-PALM), followed by a read-out of PAmCherry1-cnp1^CENP-A^ fusion using UV-activation PALM (UV-PALM). Scale bar 500 nm.

From our imaging priority (1. registering the mitotic spindle plane, 2. quantitative read-out of POI to obtain protein stoichiometry and POI cluster distribution, 3. quantitative read-out of cnp1^CENP-A^ as reference for stoichiometry read-outs and for centromere-POI-distances), literature knowledge and experience (e.g. on autofluorescence (Supplementary Figure S1) and labeling strategies (Supplementary Figure S11)), we derived several alternative measurement strategies (Supplementary Text S1). As summarized in Figure 1c, we finally decided on using dual-color Photoactivated Localization Microscopy (PALM) imaging by UV-and primed photoconversion (Turkowyd et al., 2017; Virant et al., 2017) together with a diffraction-limited SPB reference as third color. For the FP labels, we chose the bright red FP mScarlet-I (Bindels et al., 2017) as the SPB reference and the UV-photoactivatable FP PAmCherry1 (Subach et al., 2009) as the centromeric reference. For the quantitative SMLM read-out of all POIs we chose the primed photoconvertible FP mEos3.2-A69T (Turkowyd et al., 2017), whose read-out we calibrated using a FtnA 24mer standard (Supplementary Figure S8, Materials & Methods). We first created a dual-color reference strain containing the fusions for both references. Using this strain as template, we then built a triple-color (3C) strain library targeting in total 10 different key POIs of the inner and outer kinetochore (Figure 1c, Materials & Methods and Supplementary Table S1).

### Health assessment of triple-color strain library

The integration of FPs targeting important proteins of the chromosome segregation process may alter cell physiology and can cause reduced protein functionality. We performed spot tests checking for temperature sensitivity (25, 32 and 37°C) and for kinetochore-MT attachment defects using thiabendazole (2.5, 5.0, 7.5 and 10.0 mg/mL), a MT depolymerizing drug characteristic for inherent mitotic defects (Tang et al., 2013) (Supplementary Figure S2). Furthermore, we tested all cells for a wildtype-like phenotype via flow cytometry (Supplementary Figure S3). In both tests, our strains showed no deviations from the parental h^+^/WT strain except for two strains, the cnp3^CENP-C^-3C strain and the dam1-3C strain. cnp3^CENP-C^-3C exhibited both, a TBZ sensitivity in the spot tests as well as a notably larger cell size in the flow cytometry data. Due to this prominent phenotype, the cnp3^CENP-C^-3C strain was excluded from further analyses. Interestingly, cnp3^CENP-C^ is thought to interact with the N-terminal CATD region of cnp1^CENP-A^ via its C-terminus (Black et al., 2007; Carroll et al., 2010; Guse et al., 2011). Thus, the genetic tagging of both interacting protein termini might have caused the observed defects due to the proximity of the N-terminal cnp1^CENP-A^ and C-terminal cnp3^CENP-C^ FP-tags. This hypothesis is supported by measurements using a single-color strain where only the cnp3^CENP-C^ C-terminus was tagged but both reference proteins remained native. This cnp3^CENP-C^-1C strain did not show any modified phenotype (data not shown).

The dam1-3C strain appeared to be more stable in TBZ spot tests. Interestingly, similar results were measured in two other *S. pombe* studies: a deletion mutant for dam1 was hypersensitive to TBZ, whereas a C-terminal truncation mutant (dam1-127) had some higher TBZ resistance (Sanchez-Perez et al., 2005). Second, a mutant with two mutations in the C-terminal region of dam1 (H126R and E149G) also proved to be more resistant to TBZ than the wildtype (Griffiths et al., 2008). These observations suggest that the C-terminus of dam1 is important for controlling microtubule stability and that the C-terminal FP-tag in the dam1-3C strain increases stability. As the dam1-3C strain showed no mitotic delay or altered phenotype it was included in the study.

### Determining kinetochore distances and protein stoichiometry

All triple-color strain library imaging samples were prepared and imaged by a strict protocol that was repeated on different days using several biological replicates (Materials & Methods). Since single centromeres can only be resolved during metaphase/early anaphase (examples in Supplementary Figure S10, (Tournier et al., 2004)), the fraction of cells in this phase was increased via cell cycle synchronization (Supplementary Figure S2). All data was classified and postprocessed by several manual and automated SMLM data analysis steps as outlined in Supplementary Figure S5 and described in the Materials & Methods. The final, annotated data was stored in a SQL database structure. For this work, we then focused on the distances of the different POI clusters to the centromeric region marked by cnp1^CENP-A^ and their copy numbers per POI cluster. To determine the best estimates for the POI-cnp1^CENP-A^ distances, we used Bayesian inference and derived the posterior probability distribution for each POI-cnp1^CENP-A^ distance using Stan (Materials & Methods). In our Bayesian model (Supplementary Figure S6, Materials & Methods), we explicitly accounted for the angular offset between the kMT axes and the spindle axis. We visualize this offset for all cnp1^CENP-A^ centroids in Supplementary Figure S12. We further calibrated the measured localization counts per POI cluster using reference data of bacterial ferritin FtnA, an established homo-oligomer protein standard with 24 subunits (Supplementary Figure S8, Materials & Methods).

In Figure 2, we summarize all measured mean kinetochore distances in reference to cnp1^CENP-A^ and protein stoichiometries. The exact values and statistics can also be found in Tables 1 and 2. As discussed in more detail below for each POI, our *in-situ* measurements are consistent with previous work based on conventional fluorescence microscopy (these literature studies, however, show considerable variation between them) and very accurately confirm structural data from *in vitro* EM and crystallography studies. For our direct distance comparisons to structural data below, it should be considered that i) typically tens to hundreds of amino acids long sequences of the termini are unstructured and are therefore missing from structural data, which ii) also do not carry FP tags of 2-3 nm barrel size. In our results below, we will discuss the kinetochore structure from the inner kinetochore building on top of the centromeric region to the outer kinetochore proteins which facilitate microtubule attachment.

**Figure 2:**
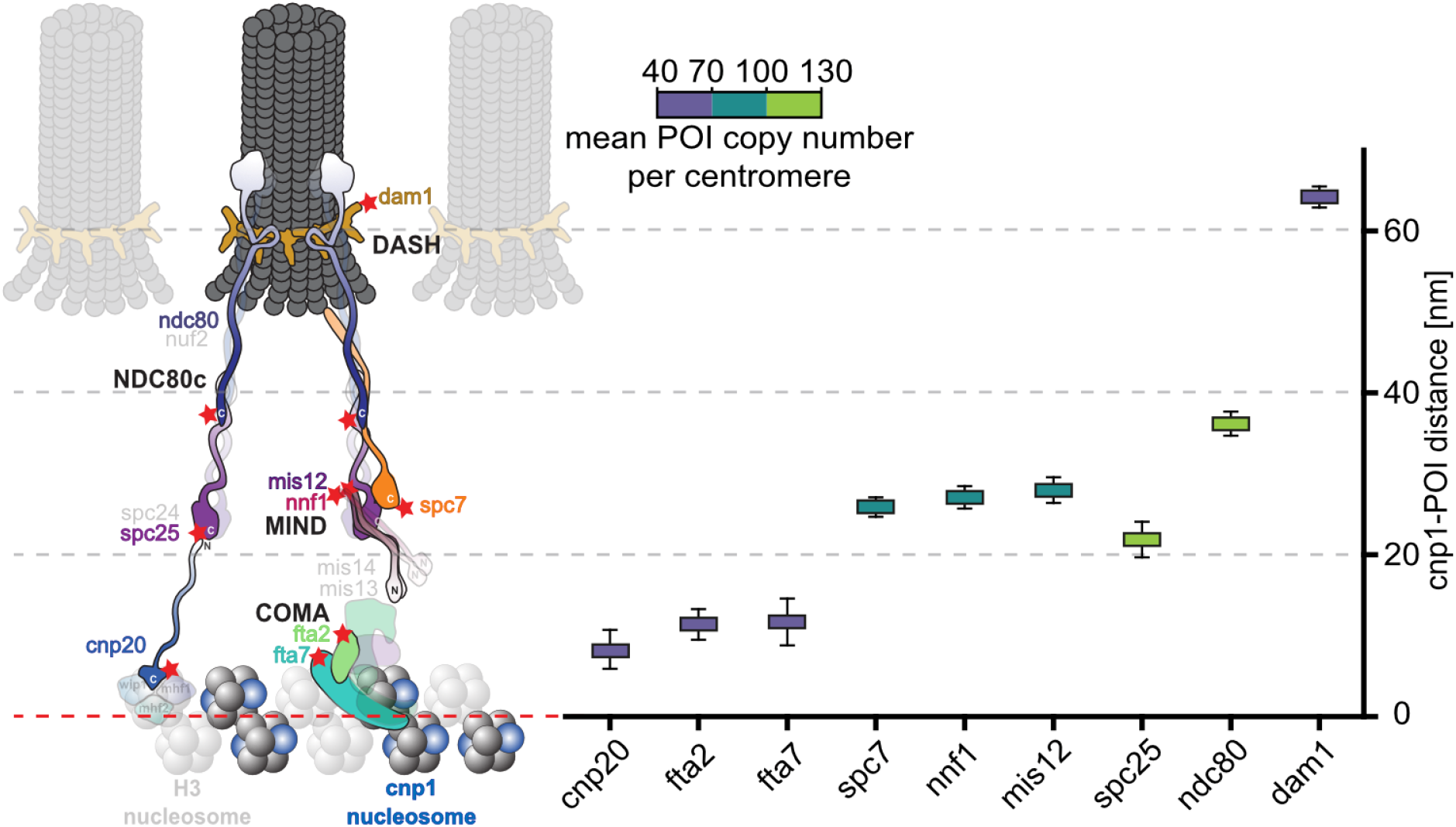
POI-cnp1^CENP-A^ distances and protein stoichiometry within the kinetochore complex. **Left** Schematic of the regional *S. pombe* centromere with parts of the inner and outer kinetochore. Whenever information was available, the shapes of POIs and subcomplexes are shown according to cryo-EM or x-ray crystallography data. Red stars mark the position of the C-terminal fluorescent protein marker mEos3.2-A69T. Structures drawn at 25% opacity were not investigated. **Right** POI distances to the reference cnp1^CENP-A^ and POI copy numbers per centromeric region as measured in this study. The colored boxes mark the mean and the whiskers the standard deviation of the posterior probability density distribution for each cnp1^CENP-A^-POI distance (Supplementary Figure S7, statistics in Table 1). The color of the box represents the mean POI copy number per cluster, as indicated by the scale bar (distributions of localization counts in Supplementary Figure S9, statistics in Table 2).

fta2^CENP-P^ and fta7^CENP-Q^ are part of the COMA complex/CCAN network and are known to possess a strict 1:1 stoichiometry with their C-termini oriented in the same direction and close to each other. Our *in situ* determined distances of 11.4 ± 1.9 nm for fta2^CENP-P^ and 11.7 ± 2.9 nm for fta7^CENP-Q^ and the POI copy numbers of 55.1 ± 3.3 and 57.4 ± 4.1 and thus in a stoichiometry of 1:1 reflect these properties and are consistent with recent cryo-EM data from the reconstituted *S. cerevisiae* COMA complex (Yan et al., 2019), where the distances from N-cnp1^CENP-A^ to fta2^CENP-P^-C and to fta7^CENP-Q^-C were determined to be 9.7 nm and 10.7 nm, respectively.

Mis12 and nnf1^PMF1^ are both part of the MIND complex (MINDc) in a 1:1 ratio (Maskell et al., 2010; Petrovic et al., 2010). Our distances are 28.0 ± 1.6 nm for mis12-C and 27.1 ± 1.4 nm for nnf1^PMF1^-C, respectively. They confirm fluorescence data where all C-termini of the MINDc proteins line up at a similar distance (Joglekar et al., 2009) and show FRET proximity signal (Aravamudhan et al., 2014) as well as EM data showing all C-termini near each other (Petrovic et al., 2014). For spc7^KNL1^, we calculate 25.9 ± 1.2 nm. Whereas full-length spc7^KNL1^ has not been purified (Maskell et al., 2010), its C-terminal structure has been resolved in EM data (Petrovic et al., 2014) and was determined to be close to the C-termini of mis12 and nnf1 (Petrovic et al., 2016). In fluorescence microscopy studies, the spc7^KNL1^ C-terminus was found to colocalize with the spc24 C-terminus of the Ndc80 complex (which is reflected in our data for spc25, see below) (Joglekar et al., 2009). Interestingly, the MINDc and spc7^KNL1^ proteins’ measured stoichiometry amounts to 1: 1.3: 1.3 (protein copy numbers of 73.1 ± 3.1 mis12: 92.6 ± 3.6 nnf1^PMF1^: 95.4 ± 3.3 spc7^KNL1^) in our hands. This differs from the reported 1: 1 ratio for mis12: nnf1^PMF1^ but posits a 1: 1 ratio for nnf1^PMF1^ and spc7^KNL1^. From biochemical reconstitution and cross-linking experiments, it is known that the MINDc has one binding site for spc7^KNL1^ at the C-terminus of mis14^NSL1^ (Petrovic et al., 2010). The sequence of that binding site (i.e. the C-termini of spc7^KNL1^ and mis14^NSL1^) is conserved over species (Petrovic et al., 2014). This places our *in situ* measured 1: 1 ratio of nnf1^PMF1^ and spc7^KNL1^ in agreement with the *in vitro* measured literature and demands for future work on mis12.

Interestingly, whereas the kinetochore structures of the outer kinetochore are highly conserved (D’Archivio and Wickstead, 2017; Meraldi et al., 2006; van Hooff et al., 2017), several strategies exist for bridging the centromere and outer kinetochore (Hamilton and Davis, 2020; van Hooff et al., 2017). Accordingly, the use and importance of the different inner kinetochore structures, i.e. either based on cnp20^CENP-T^, cnp3^CENP-C^ or fta7^CENP-Q^ (COMAc), varies between organisms (van Hooff et al., 2017): E.g. while the inner kinetochore architecture of *Drosophila melanogaster* is defined by cnp3^CENP-C^ (Ye et al., 2016), *S. cerevisiae* primarily relies on the COMAc with cnp3^CENP-C^ serving as a backup strategy (Hamilton et al., 2020; Hornung et al., 2014). Furthermore, while cnp20^CENP-T^ is non-essential in *S. cerevisiae* (Bock et al., 2012; De Wulf et al., 2003; Giaever et al., 2002; Schleiffer et al., 2012), it is the primary strategy in chicken DT40 cells (Hara et al., 2018) and human cells (Suzuki et al., 2014) which also rely on cnp3^CENP-C^ as a backup pathway (Hamilton and Davis, 2020).

To date, the preferred inner kinetochore pathway of *S. pombe* is unknown (Hamilton and Davis, 2020). The deletion of cnp3^CENP-C^ in *S. pombe* is viable (Chik et al., 2019; Kim et al., 2010; Tanaka et al., 2009) whereas deletion of cnp20^CENP-T^ is not (Tanaka et al., 2009). This is inverted for *S. cerevisiae* where mif2 (cnp3^CENP-C^) is essential but cnn1 (cnp20^CENP-T^) is not (Bock et al., 2012; De Wulf et al., 2003; Giaever et al., 2002; Kim et al., 2010; Meeks-Wagner et al., 1986; Schleiffer et al., 2012). This is also reflected in the protein copy numbers as mif2 (cnp3^CENP-C^) is more abundant than cnn1 (cnp20^CENP-T^) in S. c*erevisiae*, chicken and humans (Cieslinski et al., 2021; Johnston et al., 2010; Suzuki et al., 2015). For the COMAc, fta7 (Ame1) and mis17 (Okp1) are essential for both organisms (Hayles et al., 2013; Kim et al., 2010) and deletions of fta2 and mal2 are non-viable in *S. pombe.*

In our measurements, cnp20^CENP-T^ has a distance of 8.3 ± 2.4 nm to cnp1^CENP-A^ and thus at a similar distance as the MIND proteins fta7^CENP-Q^ and fta2^CENP-P^. Furthermore, when comparing the abundances of the inner kinetochore proteins in between the different subcomplexes, we measured a 1: 0.9 ratio of cnp20^CENP-T^ to the COMAc (Supplementary Table S4). This underlines its importance for *S. pombe* kinetochores as an essential protein (Tanaka et al., 2009) and suggests that *S. pombe* relies on the COMAc as well as the cnp20^CENP-T^ pathway in equal measures - unlike *S. cerevisiae*, which relies on the COMAc and cnp3^CENP-C^.

As representatives for the Ndc80 complex, we measured the proteins ndc80^HEC1^ and spc25 and obtained distances of 36.2 ± 1.5 nm and 21.9 ± 2.2 nm and protein copy numbers of 107.2 ± 4.3 and 119.1 ± 6.2, respectively. The shorter distance of spc25-C to cnp1^CENP-A^ in comparison to the distances of the MIND proteins to cnp1^CENP-A^ might reflect a spatial overlap of the POIs termini to interlock and stabilize the contact sites of the two sub-complexes and is in agreement with cross-linking data (Kudalkar et al., 2015). Furthermore, the order of increasing C-terminal distances from spc25 to spc7^KNL1^, nnf1^PMF1^, mis12 and ndc80^HEC1^ in our measurements correlates with decreasing FRET signal between spc25 and the POIs as measured by Aravamudhan et al. (Aravamudhan et al., 2014). A previous study shows an increase of Ndc80c and MIND complex copy numbers from metaphase to anaphase in *S. cerevisiae* and *S. pombe* cells (Dhatchinamoorthy et al., 2017). For our data, we only selected and analyzed cells in metaphase and early anaphase. Comparing the mitotic spindle length given by the sad1 spots distance and corresponding localization counts of POI and cnp1^CENP-A^ clusters in our data, we did not detect any interdependence of mitotic spindle length and POI copy numbers, which confirms our cell cycle selection (data not shown).

For the dam1/DASH complex, we calculated a distance of 64.2 ± 1.3 nm and a protein copy number of 63.7 ± 2.4 for dam1. Comparing the distance of ndc80^HEC1^-C to dam1-C, dam1 is closer than one would expect it when colocalizing with the N-term/globular head of ndc80^HEC1^ (Aravamudhan et al., 2014; Roscioli et al., 2020). Dam1 possesses a highly variable length for different fungi (Jenni and Harrison, 2018). Whereas the dam1 C-terminus has been shown to interact with the globular head of ndc80^HEC1^ in *S. cerevisiae,* it is predicted to be too short (only 155 aa in *S. pombe* in comparison to 343 aa in *S. cerevisiae*) to do so in *S. pombe* (Jenni and Harrison, 2018). Furthermore, deleting dam1 is viable for *S. pombe* (Zahedi et al., 2020) but inviable for *S. cerevisiae* (Giaever et al., 2002). Consistent with these observations, we deduct from our data that *S. pombe* dam1 indeed interacts and colocalizes with the ndc80^HEC1^-loop region. To explore this further, imaging ndc80^HEC1^ tagged at its N-terminus, within its loop region or imaging the DASH complex protein dad1 (shown to localize in the DASH complex ring structure within the ndc80^HEC1^ loop (Jenni and Harrison, 2018)) will be targeted in the future.

In contrast to the proteins of the outer kinetochore Ndc80 and MIND complexes, which exhibit higher protein abundance, our dam1 stoichiometry data shows a similarly low protein copy number as the inner kinetochore proteins from the COMAc. This was already indicated as a trend in fluorescence measurements by Joglekar et al. (Joglekar et al., 2008), where the authors measured increasing fluorescence and thus protein abundance from the inner to the outer kinetochore structure for both, *S. pombe* and *S. cerevisiae*, and a substantial drop for dam1 in *S. pombe* but not in *S. cerevisiae* cells.

## Summary

In this work, we used quantitative SMLM imaging to map the kinetochore architecture at a nanometer resolution. It is the first holistic SMLM study of a multi-protein complex in *S. pombe.* We established quantitative dual-color SMLM imaging in *S. pombe* which with we were able to drastically improve the positioning and stoichiometry accuracy of 10 kinetochore proteins being assembled *in situ* and thus within the full, native kinetochore structure. Next to drastically improving the accuracy of data of proteins that have been studied before, we add important data on proteins that a) never been quantified or b) not been quantified in *S. pombe* so far and provide nanometer-precise distances between the individual components as well as their copy numbers. This work helps to close the gap between the highly resolved *in vitro* studies and previous *in vivo* experiments. Our measurements on the structural organization of the *S. pombe* kinetochore proteins confirm the highly conserved structure of the outer kinetochore complexes between different organisms and add important details of the *S. pombe* specific outer kinetochore organization, e.g. on dam1 localization at the DASH ring. For the inner kinetochore, there exist several strategies to bridge the centromere and the outer kinetochore. Recently, increasingly more insights have been found on the differences between organisms in the use and importance of the three different inner kinetochore structures, i.e. based on either cnp20^CENP-T^, cnp3^CENP-C^ or fta7^CENP-Q^. While other studies have reported the generally different use and importance of the inner kinetochore structures between organisms, the *S. pombe* inner kinetochore architecture so far remained unexplored (Dimitrova et al., 2016; Hamilton and Davis, 2020; Petrovic et al., 2016; Przewloka et al., 2011; Screpanti et al., 2011). In our work, we were able to quantify the stoichiometry between the inner kinetochore pathways for *S. pombe*, and provide new insights into the *S. pombe* specific strategy for the inner kinetochore with an equal importance of the COMA and cnp20^CENP-T^ pathways.

The SMLM study of the *S. cerevisiae* kinetochore by Cieslinski et al. (Cieslinski et al., 2021), which was carried out in parallel to our work, is a perfect complement to our work. Together, the data consistently shows that kinetochores possess similar architecture when comparing the phylogenetically quite distant yeasts *S. pombe* and *S. cerevisiae.* The basic organizational principle of kinetochore assembly is universal despite *S. cerevisiae* maintaining a point centromere and *S. pombe* a regional structure. A detailed comparison of protein distances and stoichiometries can be found in Supplementary Table S4 and S5. Importantly, one substantial organism-specific difference for the inner kinetochore strategy surfaced: The ratio of cnp20^CENP-T^ to COMA is 1:0.9 in our case and 1:2.0 for *S. cerevisiae.* This is a strong indication that cnp20^CENP-T^ organization indeed differs between the two organisms. Furthermore, the *S. cerevisiae* work measured the position of ask1, a protein of the DASH ring. Their positioning of *S. cerevisiae* ask1 is consistent with the distance we measured for *S. pombe* dam1 and thus directly supports our reasoning that the C-terminus of *S. pombe* dam1 is localized to the DASH ring and not to the ndc80 globular heads (like for *S. cerevisiae).*

In the outlook, our strain library and results provide an excellent platform for future quantitative SMLM work on the kinetochore that goes beyond numbers and distances, and we will – by further dissecting our *S. pombe* kinetochore architecture data – next focus on structural shape details and changes during the cell cycle, as well as extend our work to medically relevant mutants.

## Supporting information

Supplementary stan code and data for distance calculations

Example csv SMLM localization data

## Author Contributions

D.L. and U.E. designed research; D.V., I.V., J.W., D.L. and U.E. designed experiments; D.V., I.V. and J.W. performed experiments; D.V., B.T., M.E. and U.E. developed analysis procedure and software; D.V., I.V., J.W., B.T., M.E. and U.E. analyzed data; D.V. and U.E. acquired funding; I.V. and J.W. designed manuscript figures and tables; U.E. wrote the paper with input from all authors.

## Acknowledgements

We are grateful to Ernest Laue for detailed discussions and critical advice and to Michael Rigl, Kai Schmidt and Bogac Aybey for their help in the laboratory, as well as to Silvia González Sierra and Victor Sourjik for their help with Flow Cytometry.

## Code availability

The Stan code and our raw data to approximate the posterior distributions of kinetochore protein distances can be found in Supplementary File 1.

