## Supplementary Figures and Text for "Unraveling the kinetochore nanostructure in *Schizosaccharomyces pombe* using multi-color single-molecule localization microscopy"

[†] authors contributed equally

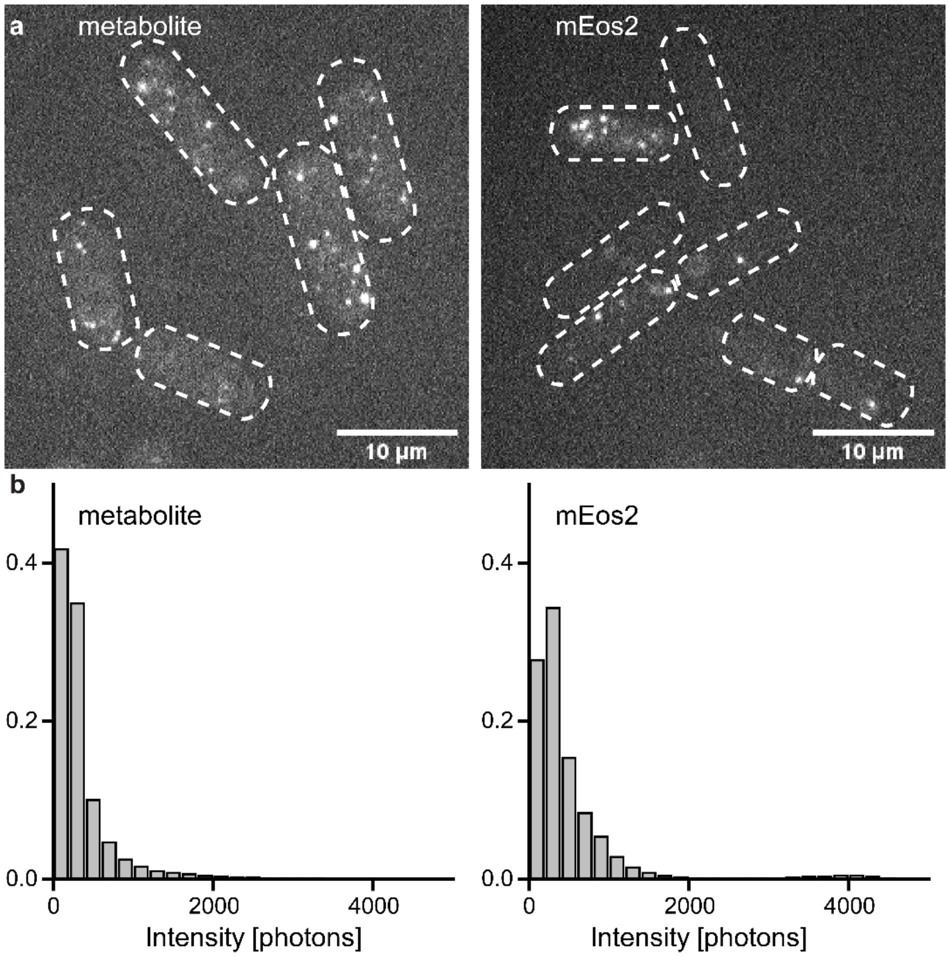


**Supplementary Figure S1: Autofluorescence of metabolites overlaps with fluorescent protein signal intensity**

**a** Exemplary images from recorded movies of *S. pombe* cells containing either accumulated metabolites (due to a mutation in the ade6 gene, left) or expressing fluorescent proteins (cytosolic mEos2, right) at similar imaging conditions. Cell borders are shown as dashed lines. Scale bar 10µm.

**b** Histograms of the intensities of individual localizations for both conditions. Localizations were filtered using a sigma of 70 - 200 nm. The metabolite signal strength overlaps with the fluorophore signal. Therefore, the autofluorescence noise cannot be reliably filtered from the fluorophore signal.

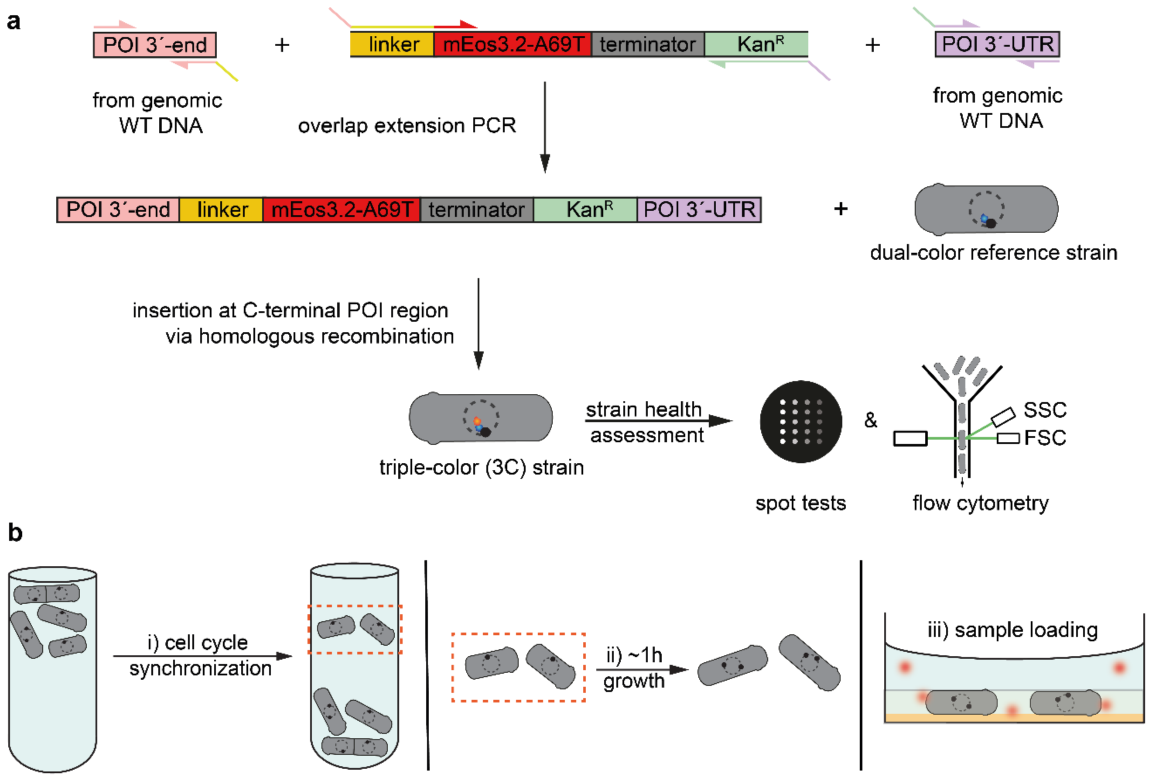


**Supplementary Figure S2: Cloning strategy and sample preparation**

**a** DNA fragments containing the POI 3'-end (pink), the FP-resistance cassette (yellow-red-grey-green) and the POI 3'- UTR (untranslated region, purple) were amplified with the corresponding primers with ~20 bp overlap to the future neighboring DNA fragments from either genomic WT DNA or a plasmid DNA. Pieces were then fused by overlap extension PCR and transformed into the dual-color reference strain (h+, leu1-32, ura4-D18, sad1:mScarlet-I:hphMX6, PAmCherry1:cnp1$^{CENP\text{-}A}$) using homologous recombination to create a triple-color strain library (Supplementary Table S1). Strain health was controlled by examining temperature and TBZ sensitivity via spot tests (Supplementary Figure S3) and normal cell length and granularity distribution via flow cytometry (Supplementary Figure S4).

**b** Cell cultures were synchronized using lactose gradient centrifugation, which accumulates cells in early G2 phase in an upper band. The cells were then extracted from the gradient column, grown for another 1-1.5 h until mitosis and chemically fixed, washed and embedded in agarose gel with fiducial markers.

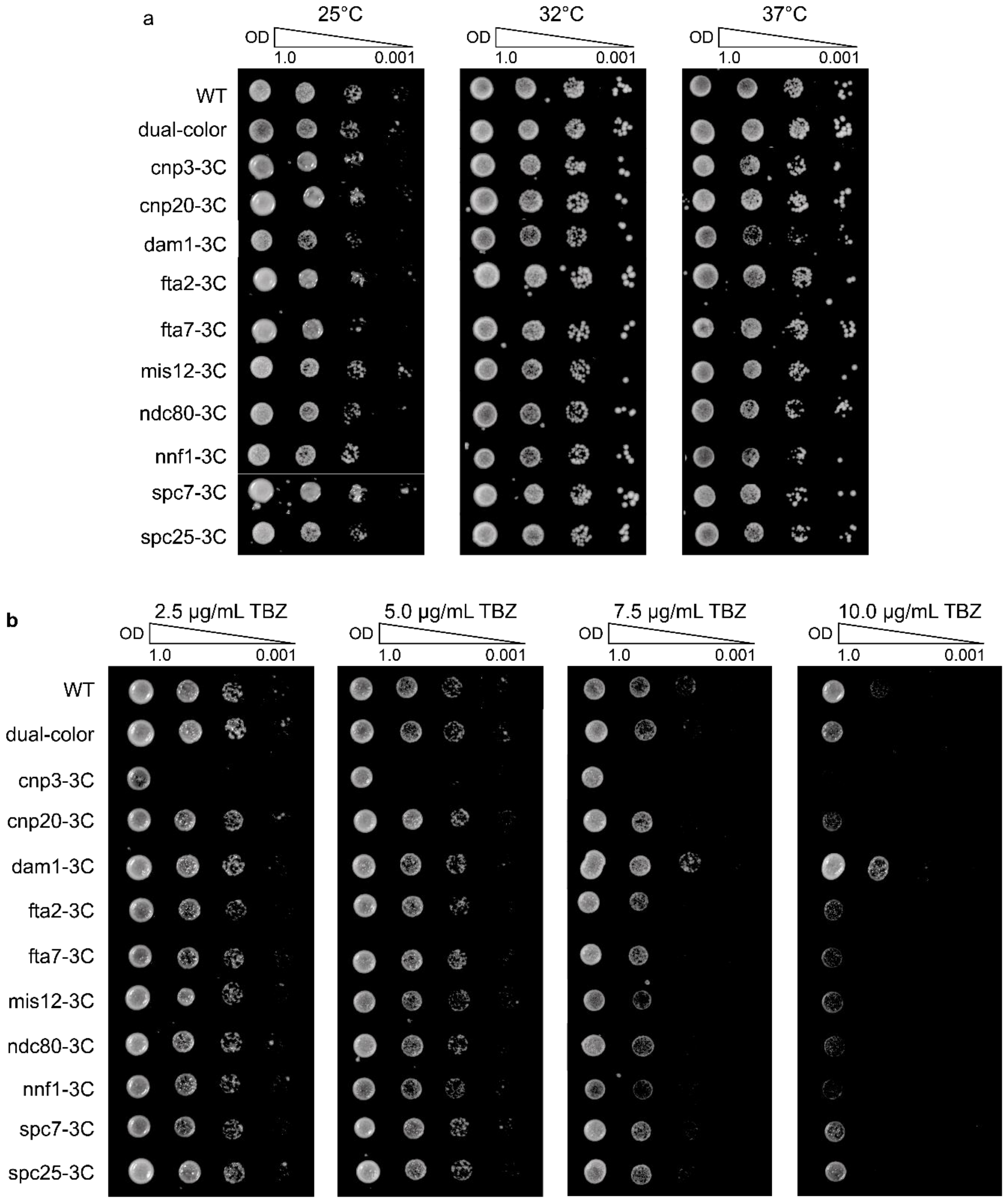


**Supplementary Figure S3: Assessment of strain health & mitotic defects**

Temperature (**a**) or thiabendazole (TBZ, which induces mitotic defects by microtubule depolymerization, **b**) sensitivities between the parental WT, the dual-color reference, and the individual triple-color (3C) strains were assessed by spot tests. Here, a tenfold dilution series of $OD_{600}$ from 1.0 to 0.001 of overnight cultures was grown on either YES media plates at 25, 32 and 37°C for three days or YES media plates containing 2.5, 5.0, 7.5 and 10.0 µg/mL TBZ and incubation at 25°C for three days.

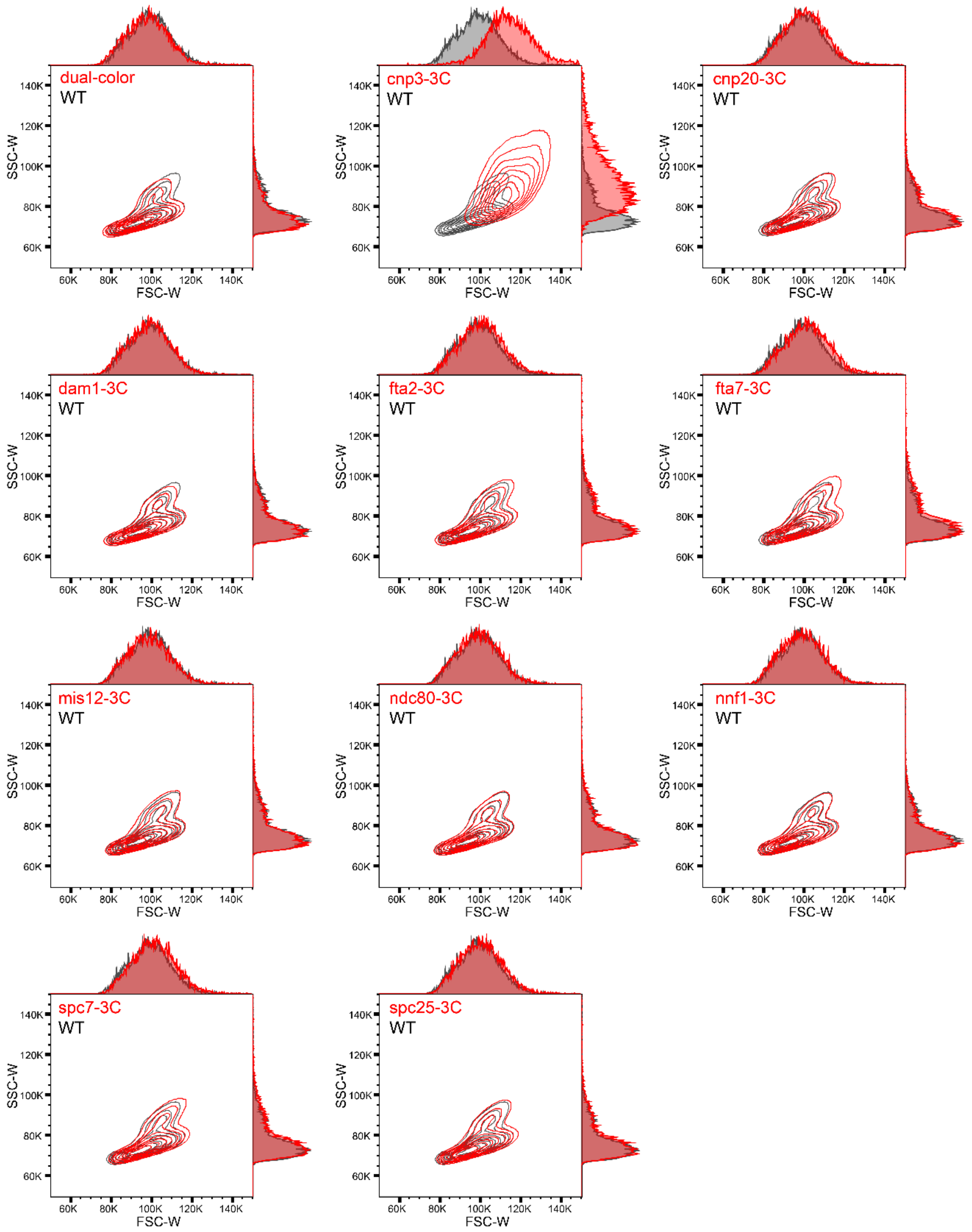


**Supplementary Figure S4: Flow cytometry measurements to assess *S. pombe* strain health**

FSC-W and SSC-W contour plots (each level within the contour plot consists of 10% measured cells) and their corresponding histograms of triple-color (POI-3C) strains and the dual-color template strain compared to a fission yeast wild-type (WT) strain. A defect in cell division usually results in increased cell length and higher FSC-W and SSC-W values, which we observed for cnp3$^{CENP\text{-}C}$-3C (top row, middle column) but none of the other strains tested.

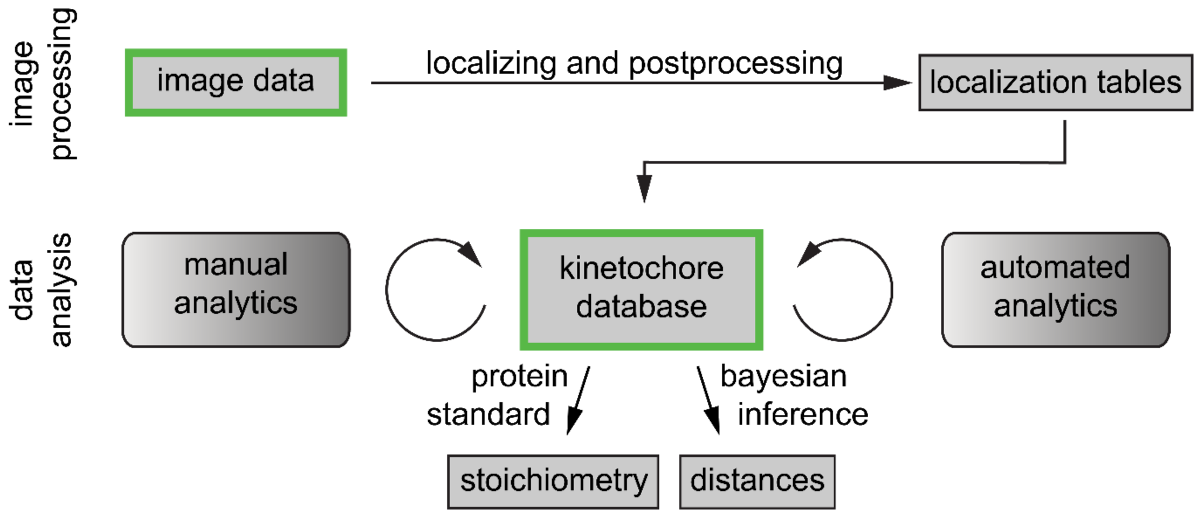


**Supplementary Figure S5: Data analysis pipeline**

Schematic representation of the data analysis pipeline. In the image processing part, image data from SMLM experiments are localized and post-processed (for quality, drift, etc.). The resulting localization tables are added to a kinetochore database, which is then used as a backend for several manual analysis (visual selection and classification steps) and automated analysis steps (channel alignment and filtering). From the database, all measures can be extracted. Here, we used localization counts per cluster and protein cluster distances to determine protein stoichiometry using a protein standard calibration and POI-cnp1$^{CENP\text{-}A}$ distances using Bayesian inference (see Materials & Methods, Supplementary Figure S6).

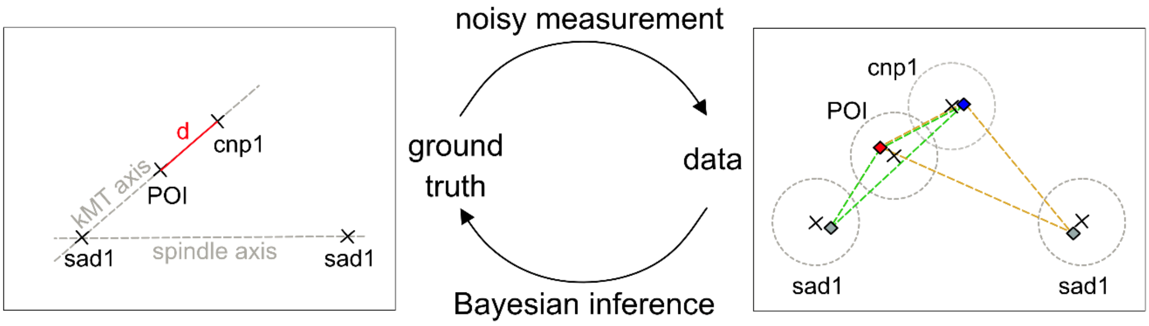


**Supplementary Figure S6: Bayesian model to estimate inner-kinetochore distances**

Schematic of our Bayesian model. To determine the real distance (marked in red) between cnp1$^{CENP\text{-}A}$ and each POI from the measured centers of their respective clusters we assumed Gaussian measurement errors of uncertain size (dotted grey circles, right). To be able to disentangle the contribution of errors and real distance in the measured data, we took the position of the associated spindle pole into account, as the centroids of sad1, POI and cnp1$^{CENP\text{-}A}$ clusters can be assumed to lie on a straight line (kinetochore microtubule (kMT) axis, left). The sad1 cluster closest to a kinetochore is not necessarily the pole to which the kinetochore is attached to. Thus, we built a mixture model to take both possibilities into account. For each kinetochore pair, we thus obtain two options to check, marked by the green and orange triangle, right.

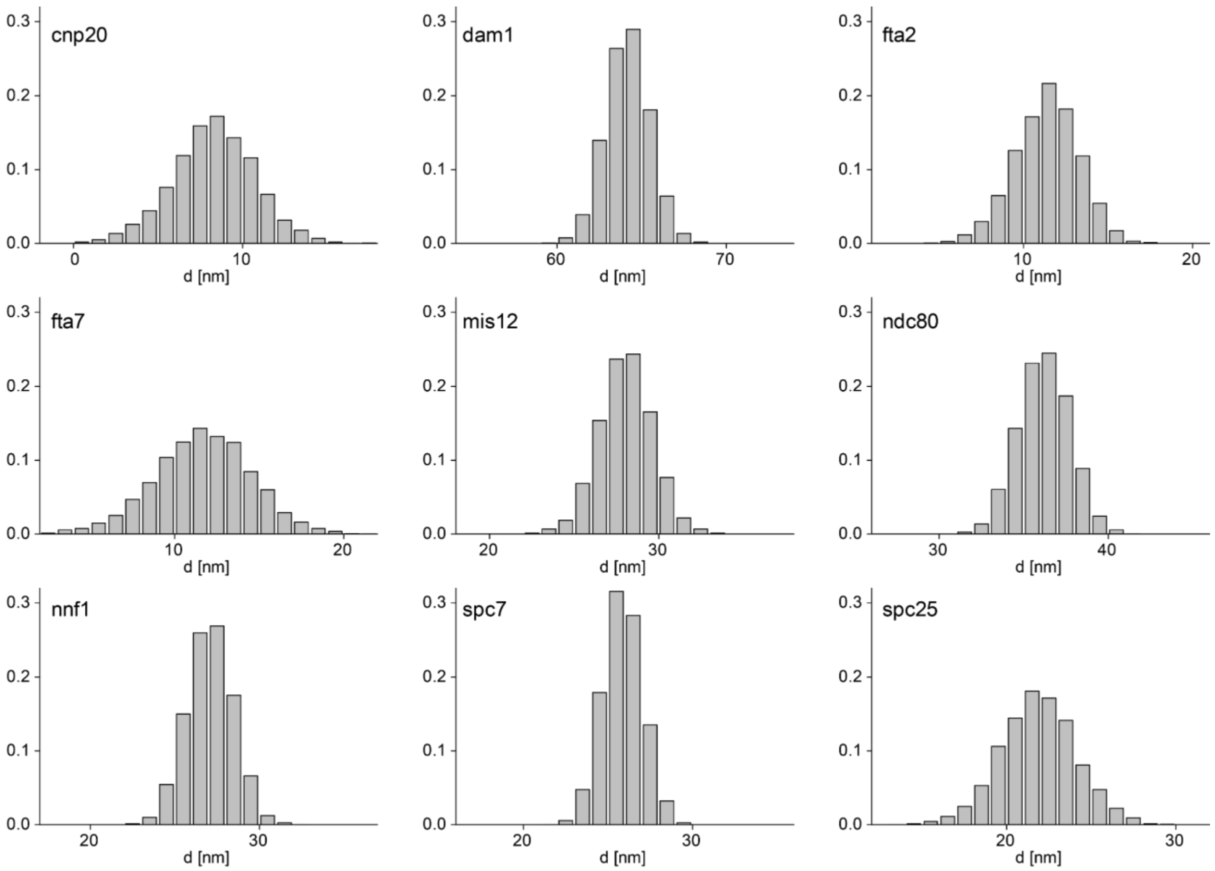


**Supplementary Figure S7: Posterior density of cnp1$^{CENP\text{-}A}$-POI distances**

The posterior density of cnp1$^{CENP\text{-}A}$-POI distances for each POI measured in this study was approximated using Hamiltonian Monte Carlo (see materials and methods). Number of centromeres used for distance measurement: N = 49 for cnp20$^{CENP\text{-}T}$, 82 for fta2$^{CENP\text{-}P}$, 58 for fta7$^{CENP\text{-}Q}$, 215 for spc7$^{KNL1}$, 161 for nnf1$^{PMF1}$, 102 for mis12, 51 for spc25, 135 for ndc80$^{HEC1}$, 155 for dam1. The code can be found in Supplementary File S1.

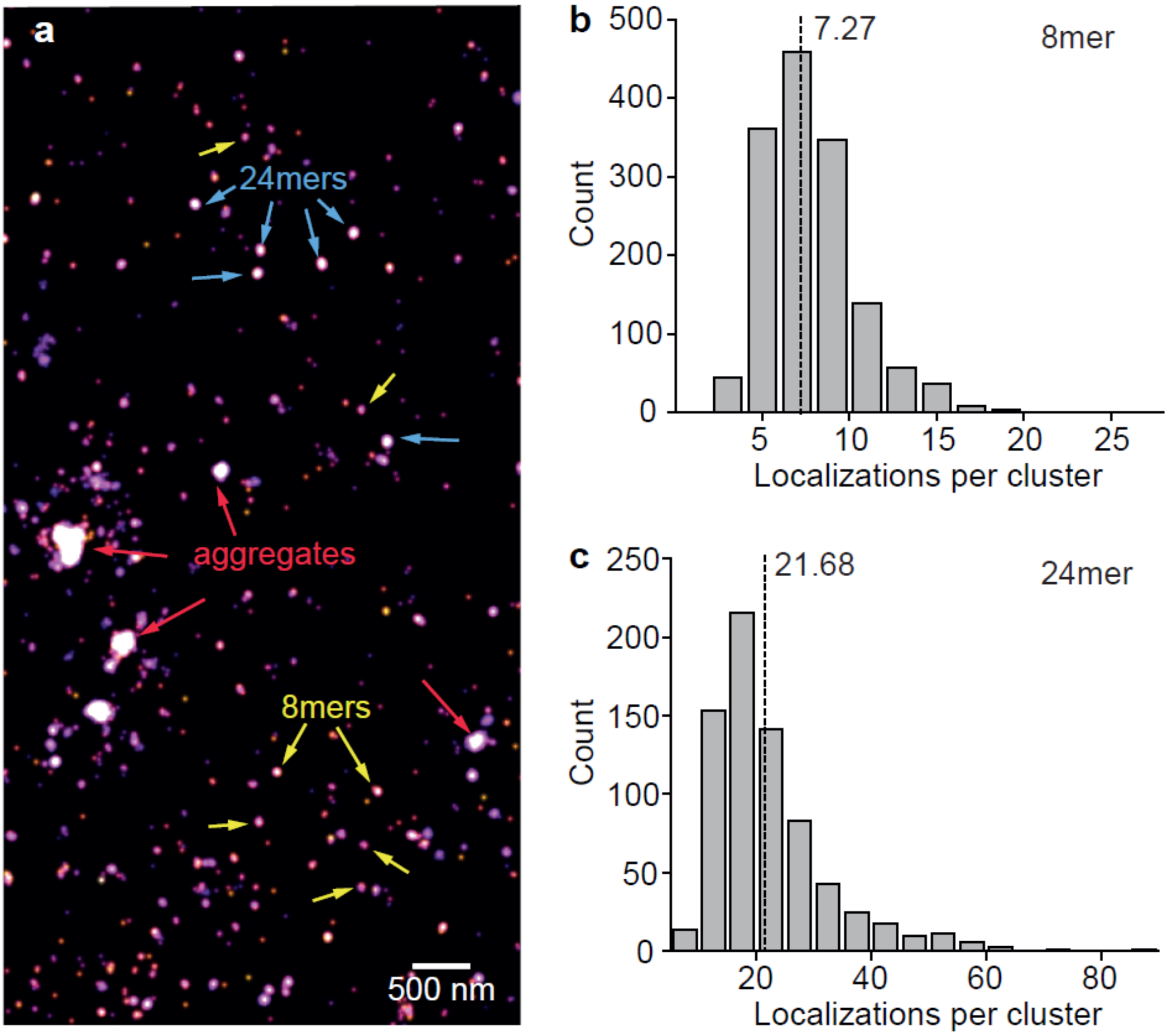


**Supplementary Figure S8: Using *E. coli* ferritin FtnA as a counting standard to calibrate POI copy numbers**

**a** Reconstructed SMLM image of isolated mEos3.2-A69T-FtnA oligomers. In our exemplary sample image, all assembly intermediates (monomers, dimers, 8mers) as well as final 24mers and some aggregates can be seen (exemplary 8mers, 24mers and aggregates are highlighted with colored arrows). Scale bar 500 nm.

**b**, **c** Histograms of localization counts per selected 8mer (b) and 24mer (c) cluster. Using the mean (dashed lines) of 7.27 ± 2.72 for 8mers and 21.68 ± 10.28 for 24mers we determine a calibration factor of 0.9. N = 1458 (8mers) and N = 725 (24mers).

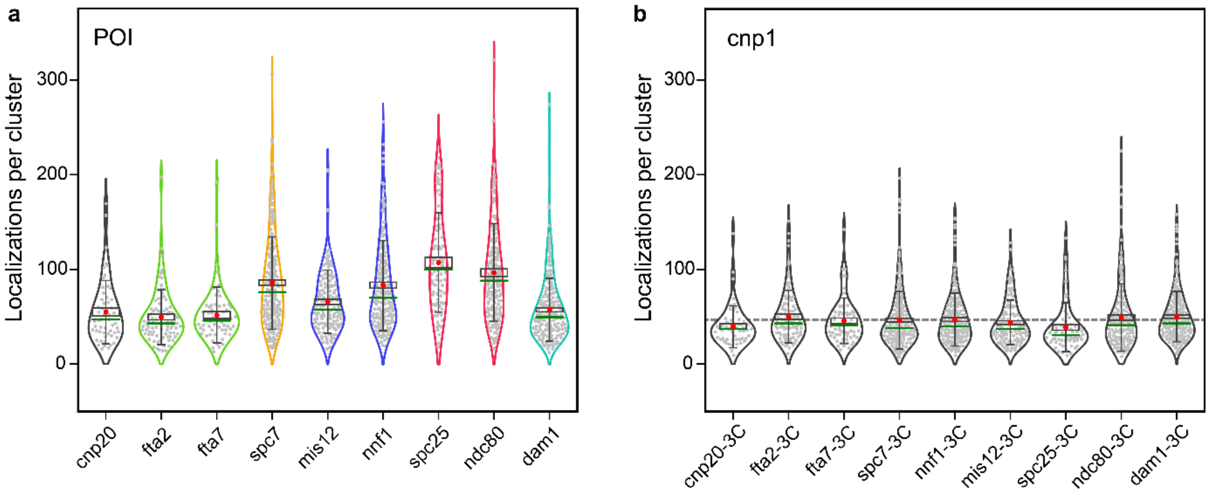


**Supplementary Figure S9: POI numbers and robustness of the method**

**a** Localizations per POI cluster. POIs from the same subcomplex are shown in the same colors (COMA (green): fta2$^{CENP\text{-}P}$, fta7$^{CENP\text{-}Q}$; MIND (blue): mis12, nnf1$^{PMF1}$; NDC80c (red): spc25, ndc80$^{HEC1}$). The red dot indicates the mean, the green line indicates the median, the black box indicates the SE of the mean, and the whiskers indicates the STD. N = 57 for cnp20$^{CENP\text{-}T}$, 87 for fta2$^{CENP\text{-}P}$, 61 for fta7$^{CENP\text{-}Q}$, 86 for spc7$^{KNL1}$, 136 for mis12, 217 for nnf1$^{PMF1}$, 86 for spc25, 172 for ndc80$^{HEC1}$, 232 for dam1.

**b** Distributions of localizations per cnp1$^{CENP\text{-}A}$ cluster are robust across different POI-3C measurements. The gray dotted line indicates the mean of all cnp1$^{CENP\text{-}A}$ clusters for reference. N = 55 for cnp20$^{CENP\text{-}T}$-3C, 96 for fta2$^{CENP\text{-}P}$-3C, 68 for fta7$^{CENP\text{-}Q}$-3C, 239 for spc7$^{KNL1}$-3C, 197 for nnf1$^{PMF1}$-3C, 122 for mis12-3C, 68 for spc25-3C, 161 for ndc80$^{HEC1}$-3C, 239 for dam1-3C.

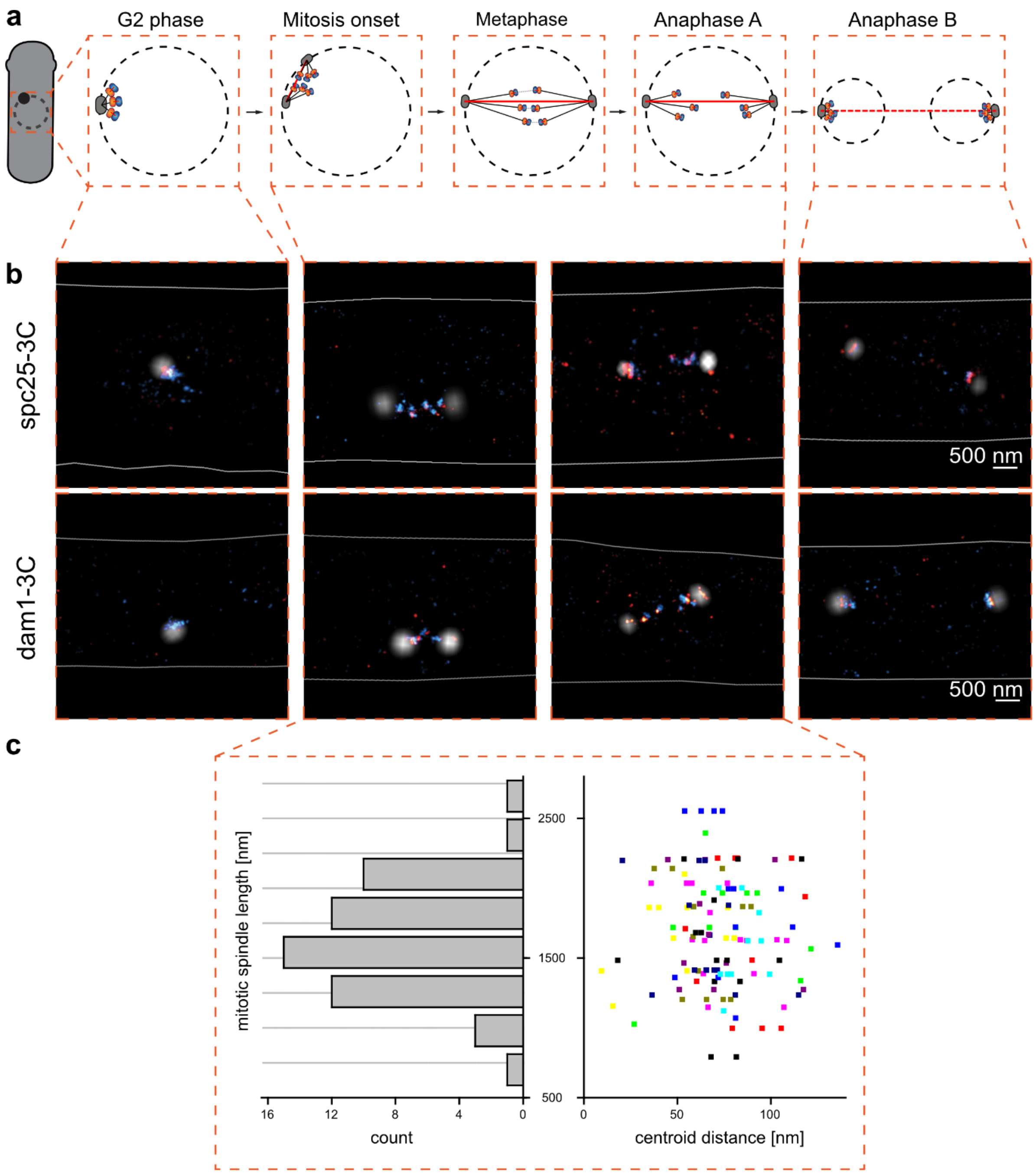


**Supplementary Figure S10: Centroid distance is independent of mitotic spindle length**

**a** Scheme of the *S. pombe* cell cycle (from left to right): A fission yeast cell in G2 phase is drawn with a nuclear envelope (NE; black dashed circle) and one spindle pole body (SPB, black). Insets show the nucleus changing over the cell cycle phases (G2, onset of mitosis, metaphase, Anaphase A and Anaphase B) with the kinetochore (orange) linking the centromere (blue) to the SPB via a bundle of kinetochore microtubules (black lines), while the spindle microtubules (red line) push the two SPBs further apart.

**b** Exemplary three-color SMLM images of spc25-3C (top row) and dam1-3C (bottom row) strains representing the cell cycle stages shown in (**a**). SPB (sad1-mScarlet-I) localizations are shown in white, kinetochores (POI-mEos3.2-A69T) in red, centromeres (PAmCherry1-cnp1$^{CENP\text{-}A}$) in blue, and the cell border (determined by the bright light image) is drawn as a white line. Scale bar 500 nm.

**c** Left: Histogram of an exemplary data set of mitotic spindle lengths (distance between the two SPBs during mitosis) for the POI dam1. Right: the distance between the centroids of individual kinetochore cluster pairs (POI and cnp1$^{CENP\text{-}A}$) plotted against the mitotic spindle length from the same cell. Data points of the same height and color are from the same mitotic spindle. All spindle lengths are shorter than the average nuclear diameter of 2-3 µm, thus excluding anaphase B cells.

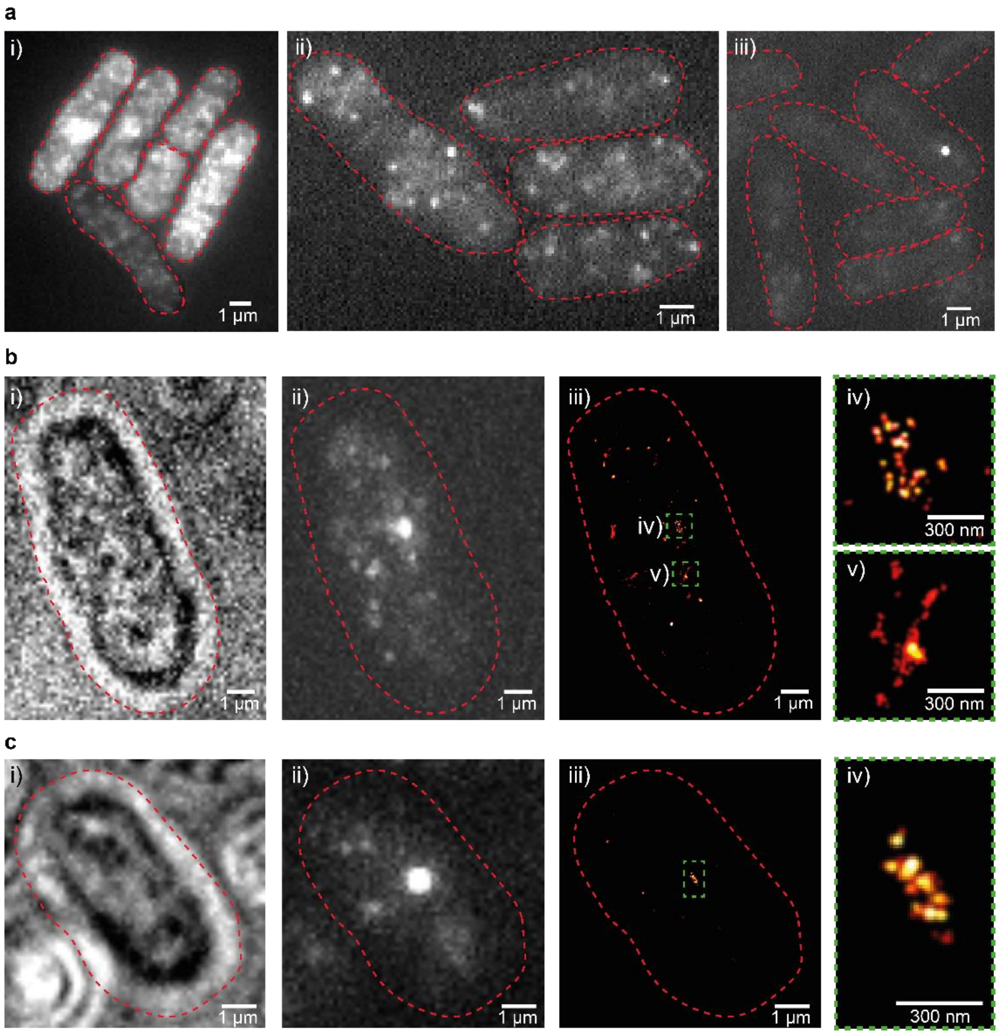


**Supplementary Figure S11: Dye staining in *S. pombe* is heterogeneous and unreliable for low copy number targets**

**a** Staining of Halo-cnp1$^{CENP\text{-}A}$ cells with the cyanine Alexa Fluor 647 (i) 200 nM, ii) 50 nM) shows a high degree of nonspecific labeling, even when using the blocking agents BSA and Image-IT (which reduce nonspecific interactions of the charged dye with the sample). Staining with iii) 200 nM of cyanine CF680 shows low labelling efficiency even after cell wall and membrane were partially digested with zymolyase and Triton-X100. We attribute the low efficiency to its large molecular weight (CF680 is about 2.3 times larger than Alexa Fluor 680 as it has masking groups to avoid nonspecific staining due to charges).

**b** Exemplary cell showing high unspecific staining of CF647-Halo-cnp1$^{CENP\text{-}A}$ in i) brightlight, ii) conventional fluorescence and iii) SMLM imaging. Detailed views illustrating iv) the cnp1$^{CENP\text{-}A}$ signal compared to some v) nonspecific signal. The iv) signal also shows a low labeling efficiency with low localization counts (compared to the cell in **b**)

**c** Exemplary cell with low, nonspecific staining of CF647-Halo-cnp1$^{CENP\text{-}A}$ at a high labeling efficiency.

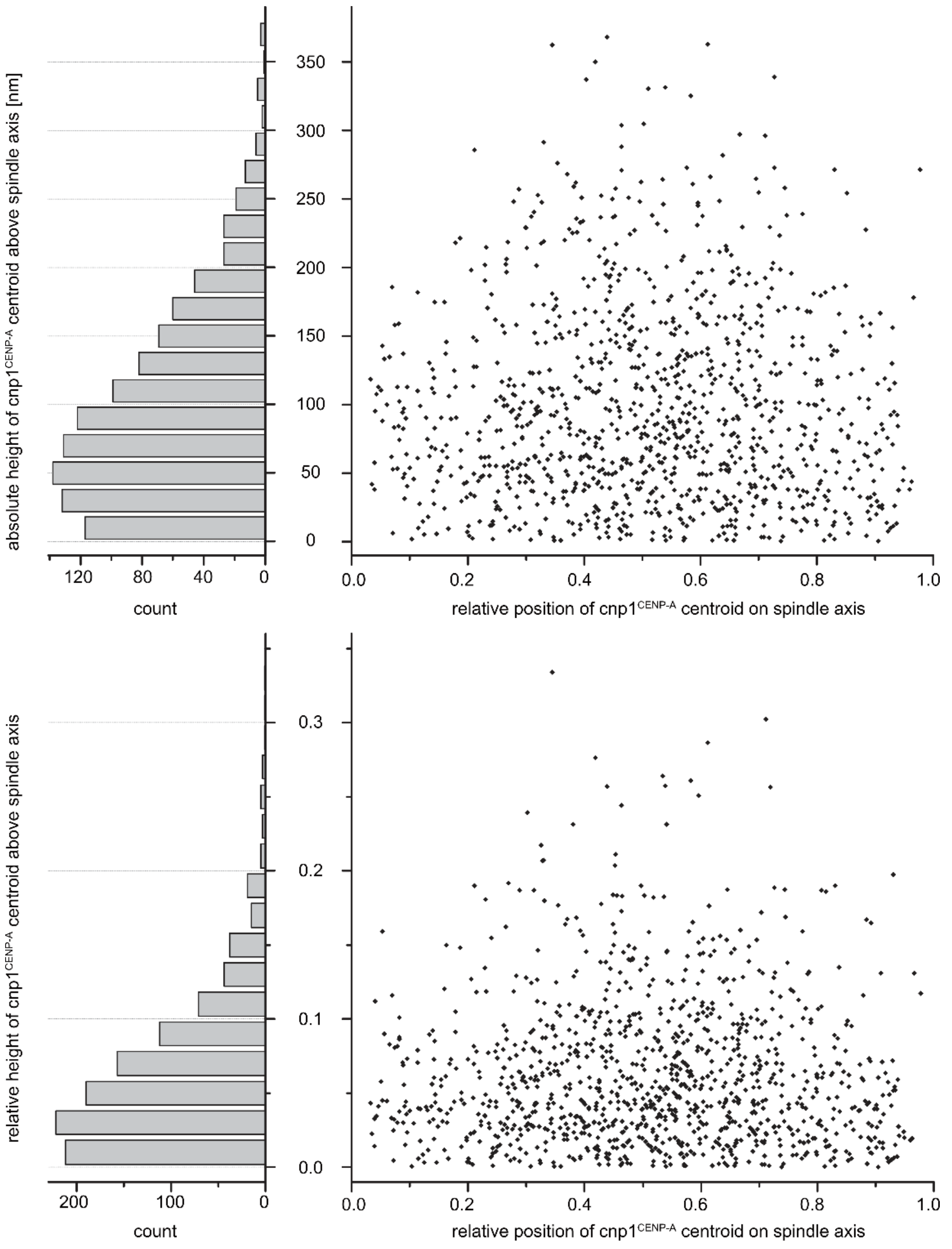


**Supplementary Figure S12: Angular offset of kMT and spindle axes**

Plotted are relative position and height over the spindle axis (defined as sad1-sad1 centroid distance) for all measured cnp1$^{CENP-A}$ centroids. Height of cnp1$^{CENP-A}$ centroids is either plotted in absolute nanometer distances (top) to visualize that most kinetochores are in direct vicinity to the central bundle or normalized to the respective spindle length of the cells (bottom) to represent the angular distribution between the spindle and kMT axes.

| Strain | Genotype | Source |
|---|---|---|
| SP176 (972) | wt h- | gift from Laue lab, Cambridge, UK |
| SP177/ wt h+ | h+, ade6-210, leu1-32, ura4-D18 | (Lando et al., 2012) |
| SP11/ DL70 | h+, ade6-210, leu1-32, ura4-D18, PAmCherry1:cnp1 | (Lando et al., 2012) |
| SP118 | h-, sad1:mScarlet-I:hphMX6 | this study |
| SP145 | h+, leu1-32, ura4-D18, sad1:mScarlet-I:hphMX6, PAmCherry1:cnp1 | this study |
| SP137 | h+, leu1-32, ura4-D18, spc25:mEos3.2-A69T:kanMX6, sad1:mScarlet-I:hphMX6, PAmCherry1:cnp1 | this study |
| SP141 | h+, leu1-32, ura4-D18, mis12:mEos3.2-A69T:kanMX6, sad1:mScarlet-I:hphMX6, PAmCherry1:cnp1 | this study |
| SP144 | h+, leu1-32, ura4-D18, dam1:mEos3.2-A69T:kanMX6, sad1:mScarlet-I:hphMX6, PAmCherry1:cnp1 | this study |
| SP146 | h+, leu1-32, ura4-D18, fta2:mEos3.2-A69T:kanMX6, sad1:mScarlet-I:hphMX6, PAmCherry1:cnp1 | this study |
| SP147 | h+, leu1-32, ura4-D18, fta7:mEos3.2-A69T:kanMX6, sad1:mScarlet-I:hphMX6, PAmCherry1:cnp1 | this study |
| SP150 | h+, leu1-32, ura4-D18, cnp3:mEos3.2-A69T:kanMX6, sad1:mScarlet-I:hphMX6, PAmCherry1:cnp1 | this study |
| SP152 | h+, leu1-32, ura4-D18, ndc80:mEos3.2-A69T:kanMX6, sad1:mScarlet-I:hphMX6, PAmCherry1:cnp1 | this study |
| SP153 | h+, leu1-32, ura4-D18, nnf1:mEos3.2-A69T:kanMX6, sad1:mScarlet-I:hphMX6, PAmCherry1:cnp1 | this study |
| SP154 | h+, leu1-32, ura4-D18, spc7:mEos3.2-A69T:kanMX6, sad1:mScarlet-I:hphMX6, PAmCherry1:cnp1 | this study |
| SP155 | h+, leu1-32, ura4-D18, cnp20:mEos3.2-A69T:kanMX6, sad1:mScarlet-I:hphMX6, PAmCherry1:cnp1 | this study |
| SP109 | h+, ade6-210, leu1-32, ura4-D18, Halo:cnp1 | (Vojnovic, 2016) |
| SP16 | h+, ade6-210, leu1-32, ura4-D18, pREPnmt81-mEos2::Leu2 | (Lando et al., 2012) |
| EC290 | Rosetta DE3 pRSETa mEos3.2-A69T:FtnA | this study |

**Supplementary Table S1: *S. pombe* and *E. coli* strains used in this study**

| Primer name | 5’→3’ SEQUENCE + overlap |
|---|---|
| Ade6_F2 | CTCATTAAGCTGAGCTGCCAAG |
| Ade6_R2 | TGCATAGGCGACCATAGACAT |
| AGGSG_mEos3.2_F | gccggaggcagtggttct |
| Cnp3_F1 | GGAAAGATCGAGGTCACAGT |
| Cnp3_F2 | aatgctggtcgctatactgctgtcCAATACTAATAGTGTGTTATGGATTTCG |
| Cnp3_R1_AGGSG | GGGATTTTCCAAACGAACGAgccggaggcagtggt |
| Cnp3_R2 | AAAGTCAAATCTAACGGTCGC |
| Cnp20_F1 | TTCAACACTTGTGCTACCGGAAA |
| Cnp20_F2 | aatgctggtcgctatactgctgtcTGCGTACTTCTCCTTTACATTCATC |
| Cnp20_R1_AGGSG | ACCTCCGGCAATTAAGAGAACCgccggaggcagtggt |
| Cnp20_R2 | CGCTTGATTCGATACACTTACAAGT |
| Dam1-GFP F1 | TGCCGAAAGCGCTGTAGAA |
| Dam1F2 | aatgctggtcgctatactgctgtcATTTATTTAAGCAAGGGAGACTGGTTG |
| Dam1_R1_Overhang_ AGGSG_D2 | AGAAACCTATTCCGCTTCCAGAgccggaggcagtggttct |
| Dam1R2 | TAGCTTCTCCAATCTTCAATTTCCA |
| fta2_F1 | CATGGACGCTCAATGTTTCT |
| fta2_F2 | aatgctggtcgctatactgctgtcGAAGGATAAATTGATATTTTTAACATGGTT |
| fta2_R1_AGGSG | GGCATTATTTAAACCTCATTTGGGGgccggaggcagtggt |
| fta2_R2 | AGTTCTTTTGGCAGAATGGG |
| Fta7_F1 | TTCAGACTCCAACGATTTCTC |
| Fta7_F2 | aatgctggtcgctatactgctgtcACATAGAAAAGCTAGAGCTTAAGAC |
| Fta7_R1_AGGSG | AGTCATCCAAACTTAAGATAAAGAATATCgccggaggcagtggt |
| Fta7_R2 | GAGTTTAGGGTAGGGTAAGCA |
| KanR_cassette_R | Aatgctggtcgctatactgctgtc |
| Mis12_F1 | TCTGCAGCATGCCGTTAAAAG |
| Mis12_F2 | aatgctggtcgctatactgctgtcTACTAATCAACTAGCTAAAGTCTTGAGATG |
| Mis12_R1_AGGSG | CGGACATACTGACGAGCCTgccggaggcagtggt |
| Mis12_R2 | TCTGACCCATTAACTCCAAATCTGT |
| Ndc80_F1 | ACAACAGCTCAAACTTTCTTCG |

| | |
|---|---|
| Ndc80_F2 | aatgctggtcgctatactgctgtcATTCTATTCATCGTATTGTGCTGTC |
| Ndc80_R1_AGGSG | ACCTATCTCGTTCGGAACTGgccggaggcagtggt |
| Ndc80_R2 | ACGAACTGTTTTGGCTAAAAATTTG |
| Nnf1_F1 | AAGCTTAATCAGGATCTGTTGG |
| Nnf1_F2 | aatgctggtcgctatactgctgtcAAGAAGTAAATTTCTAATCAGTTGCA |
| Nnf1_R1_AGGSG | AACGAACAAGGAAATATAGAACGTgccggaggcagtggt |
| Nnf1_R2 | CTCAAAAACATTCCAACGCAA |
| Spc7_F1 | TTGAGCTATACCTGCGTTCGG |
| Spc7_F2 | aatgctggtcgctatactgctgtcATATTAATGGGAATGATTAGCTATGCTGC |
| Spc7_R1_AGGSG | CTTGTCTTACTGTTTGGAACAAATACAGCgccggaggcagtggt |
| Spc7_R2 | AAACCCGTAATGCGCTACAAAA |
| Spc25_F1 | ATCAATCTTGCTGAAAGGGATTA |
| Spc25_F2 | aatgctggtcgctatactgctgtcAGATCTTCTTCTTGTTTAACATAAACTT |
| Spc25_R1_AGGSG | TAGAAAGGATCTGTCTCAATTGATTgccggaggcagtggt |
| Spc25_R1_D2 | TAGAAAGGATCTGTCTCAATTGATTgccggaggcagtggtATG |
| Spc25_R2 | AACTGGTGGAATCCATGGT |
| mEos3.2_F_pRSET | ACGATAAGGATCGATGGGGATCCATGtctgccattaaaccgga |
| mEos3.2_R_FtnA | ggatgctccgctagccttgcgacgcgcattatcc |
| pRSET_F_preFtnA_68 | aaggctagcggagcatcc |
| pRSET_R_blunt_68 | GGATCCCCATCGATCCTTATCGT |

**Supplementary Table S2: List of primers used for constructing the 3C-library and the FtnA protein standard as listed in table S1**

| complex | components | expected ratios | measured ratios | weighted mean | STD |
|---|---|---|---|---|---|
| **COMAc** | fta2, fta7, mal2, mis17 | 1:1:1:1 | 1:1.0:NA:NA | 56.0 | 28.2 |
| **MINDc** | mis12, nnf1, mis13, mis14 | 1:1:1:1 | 1:1.3:NA:NA | 85.1 | 43.1 |
| **NDC80c** | ndc80, nuf2, spc24, spc25 | 1:1:1:1 | 1:NA:NA:1.1 | 111.2 | 51.7 |

**Supplementary Table S3: Weighted mean of localization counts for POIs belonging to the same kinetochore subcomplex**

| complexes | ratio | reference | host | method | cell cycle stage |
|---|---|---|---|---|---|
| **cnp20$^{CENP-T}$ : COMA** | 1 : 0.9 | this study | *S. pombe* | SMLM imaging | Meta- to Anaphase A |
| | 1: 2.0 | (Cieslinski et al., 2021) | *S. cerevisiae* | SMLM imaging | Metaphase |
| **MIND : spc7$^{KNL1}$** | 1 : 1.1 | this study | *S. pombe* | SMLM imaging | Meta- to Anaphase A |
| | 1 : 0.7 | (Joglekar et al., 2008) | *S. pombe* | fluorescence ratio | G2 to Metaphase |
| | 1 : 1.0 | (Joglekar et al., 2008) | *S. pombe* | fluorescence ratio | Anaphase to Telophase |
| | 1 : 0.7 | (Lawrimore et al., 2011) | *S. pombe* | corrected from Joglekar et al. 2008 | G2 to Metaphase |
| | 1 : 1.3 | (Joglekar et al., 2006) | *S. cerevisiae* | fluorescence ratio | Metaphase |
| | 1 : 1.1 | (Joglekar et al., 2006) | *S. cerevisiae* | fluorescence ratio | Anaphase |
| | 1 : 1.0 | (Lawrimore et al., 2011) | *S. cerevisiae* | fluorescence ratio | Anaphase |
| | 1 : 0.7 | (Dhatchinamoorthy et al., 2017) | *S. cerevisiae* | fluorescence ratio | Anaphase |
| | 1 : 1.1 | (Cieslinski et al., 2021) | *S. cerevisiae* | SMLM imaging | Metaphase |
| | 1 : 0.8 | (Johnston et al., 2010) | *Chicken DT40* | fluorescence ratio | Metaphase |
| | 1 : 0.8 | (Lawrimore et al., 2011) | *Chicken DT40* | fluorescence ratio | Metaphase |
| | 1 : 0.8 | (Emanuele et al., 2005) | *X. laevis* | Biochemical assay | unsynchronized |
| **COMA : MIND** | 1 : 1.5 | this study | *S. pombe* | SMLM imaging | Meta- to Anaphase A |
| | 1 : 1.6 | (Joglekar et al., 2008) | *S. pombe* | fluorescence ratio | G2 to Metaphase |
| | 1 : 0.8 | (Joglekar et al., 2008) | *S. pombe* | fluorescence ratio | Anaphase to Telophase |
| | 1 : 1.6 | (Lawrimore et al., 2011) | *S. pombe* | corrected from Joglekar et al. 2008 | G2 to Metaphase |
| | 1 : 2.2 | (Joglekar et al., 2006) | *S. cerevisiae* | fluorescence ratio | Metaphase |
| | 1 : 2.3 | (Joglekar et al., 2006) | *S. cerevisiae* | fluorescence ratio | Anaphase |
| | 1 : 2.3 | (Lawrimore et al., 2011) | *S. cerevisiae* | fluorescence ratio | Anaphase |
| | 1 : 2.3 | (Dhatchinamoorthy et al., 2017) | *S. cerevisiae* | fluorescence ratio | Anaphase |
| | 1 : 1.8 | (Cieslinski et al., 2021) | *S. cerevisiae* | SMLM imaging | Metaphase |

**Supplementary Table S4: POI copy number ratios between different kinetochore subcomplexes**

| | Kinetochore subcomplex | POI (homolog in *S. cerevisiae*) | distance [nm] ± STD to *S.pombe* C-term spc7$^{KNL1}$ (spc105), this study | distance [nm] ± SEM to *S. cerevisiae* C-term spc7$^{KNL1}$ (spc105), Cieslinski et al. |
|---|---|---|---|---|
| **cnp1$^{CENP\text{-}A}$** | | **cnp1$^{CENP\text{-}A}$ (cse4)** | N-term: -25.9 ± 1.2 | C-term: -16.9 ± 1.3 |
| **CCAN** | **CBF3** (only in cerevisiae) | **N/A (cep3)** | / | -21.1 ± 1.7 |
| | | **cnp3 (mif2)** | / | -23.8 ± 2.0 |
| | **CENP-T/cnn1** | **cnp20$^{CENP\text{-}T}$ (cnn1)** | -17.6 ± 2.7 | -20.1 ± 2.7 |
| | **COMA** | **fta2$^{CENP\text{-}P}$ (ctf19)** | -14.5 ± 2.2 | -14.9 ± 1.7 |
| | | **fta7$^{CENP\text{-}Q}$ (okp1)** | -14.2 ± 3.1 | -13.4 ± 1.4 |
| | **CENP-N/Chl4** | **mis15 (chl4)** | / | -23.5 ± 2.9 |
| **KMN** | **KNL1/Spc105** | **spc7$^{KNL1}$ (spc105)** | 0 | 0 |
| | **MIND** | **nnf1$^{PMF1}$** | 1.2 ± 1.8 | 4.8 ± 2.6 |
| | | **mis12 (mtw1)** | 2.1 ± 2 | 4.3 ± 0.6 |
| | | **mis13 (dsn1)** | / | 3.1 ± 0.6 |
| | | **mis14 (nsl1)** | / | 6.5 ± 1.5 |
| | **NDC80** | **spc25** | -4 ± 2.5 | -2.5 ± 0.8 |
| | | **ndc80$^{HEC1}$** | 10.3 ± 1.9 | 13.6 ± 1.2 |
| | | **nuf2** | / | 16.9 ± 1.5 |
| **Dam1/DASH** | **Dam1/DASH** | **dam1** | 38.3 ± 1.8 | / |
| | | **ask1** | / | 44.3 ± 1.8 |

**Supplementary Table S5:** Comparison of protein cluster distances between our study in *S. pombe* and the study of Cieslinski et al. in *S. cerevisiae*. The studies used different reference proteins. Whereas our studies used N-terminal cnp1$^{CENP\text{-}A}$ as the reference, Cieslinski et al used C-terminal spc7$^{KNL1}$. In the upper table, we thus converted our numbers and errors into the Cieslinski et al. reference frame by using our distance between C-terminal spc7$^{KNL1}$ and N-terminal cnp1$^{CENP\text{-}A}$. From the literature, it is known that the intramolecular C- to N-terminal cnp1$^{CENP\text{-}A}$ distance is about 3 - 5 nm (Migl et al., 2020; Sekulic et al., 2010; Tachiwana et al., 2011; Yan et al., 2019) which should be considered when comparing the numbers in the above table.

### Supplementary Text S1

#### Choosing a labeling strategy for quantitative SMLM in *S. pombe*

While the number of SRM studies in fission yeast being published has been increasing in recent years, the majority of them are single color studies that target individual proteins (Akamatsu et al., 2017; Bell et al., 2014; Etheridge et al., 2014; Etheridge et al., 2021; Lando et al., 2012; Laplante et al., 2016; Matsuda et al., 2015), and only a few are dual-color studies visualizing several targets at the same time (Bestul et al., 2021; McDonald et al., 2017; Virant et al., 2017). Among those studies, single-molecule localization microscopy (SMLM) takes on a special role, as this technique does not only offer increased resolution, but also allows the determination of protein stoichiometries (Akamatsu et al., 2017; Lando et al., 2012; Laplante et al., 2016). A detailed review highlighting multi-color single-molecule localization microscopy studies in microbial organisms can be found in Vojnovic et al. (Vojnovic et al., 2019).

To construct a nanoscale map of the kinetochore in *S. pombe*, we labeled three proteins, the protein of interest, the centromeric reference cnp1$^{\text{CENP-A}}$ and the spindle pole protein sad1. Imaging the spindle poles enabled us to a) easily identify and read out the correct cell cycle stage and focal plane and to b) drastically improve the accuracy of our distance calculations. The latter is possible as all three protein(-clusters) can be assumed to lie on a straight line, which allows a direct estimation of the size of the measurement errors from the deviation of that line (see data analysis and Supplementary File S1).

As all organisms and targets have different requirements and specifics, SMLM labeling strategies, which need to be highly optimized to produce satisfying results, must be specifically adapted and tested for the actualities of each system (Vojnovic et al., 2019). In *S. pombe*, we avoided the green and blue color channels, due to increased phototoxicity as well as the fact that we found high levels of autofluorescence in those channels, as *S. pombe* is easily stressed by unstable or insufficient pH, temperature or aeration.

Furthermore, we found a high level of autofluorescence due to a selection marker in standard laboratory *S. pombe* strains which works by interrupting the adenine biosynthesis pathway to accumulate a precursor molecule – a bright red pigment which can be easily seen in screening colonies (Allshire, 1995; Levenberg and Buchanan, 1957; Lukens and Buchanan, 1959). These adenine deficient laboratory strains (most common are ade6-M210 and ade6-M216), even when grown under full adenine supply, possess a background level of metabolites which strongly disturb sensitive SMLM measurements even when colonies or liquid cultures remain inconspicuously colorless (Supplementary Figure S1 and (Winkelmeier, 2018)). We thus "cured" the adenine gene using a wildtype strain template for all strains in this study.

For three labels we were in need for three fluorophores with mutually supporting SMLM imaging properties: In general, *in situ* targeting of proteins can be achieved extrinsically by using dyes (e.g. using immunofluorescence and enzyme or peptide tags) or intrinsically by using FPs as genetic tags. On the one hand, organic dyes are brighter and therefore have a higher signal-to-noise ratio and localization precision, but on the other hand they bring two disadvantages: a) extrinsic staining is accompanied by non-specific staining and has unknown and possibly even heterogeneous staining efficiency, and b) SMLM imaging of organic dyes requires imaging buffers and might exhibit heterogeneous switching behavior. These drawbacks significantly affect *any* quantitative SMLM study but become especially severe for low copy number targets and for multi-color studies. While evaluating different labeling and imaging strategies, we tested different dyes. Surprisingly, the "golden SMLM standard" Alexa Fluor 647 does not work reliably in *S. pombe* (highly heterogeneous blinking to even non-blinking behavior - which we not only found for live cells but also for fixed cells and for all buffers tested (see buffer table in (Turkowyd et al., 2016)) for reasons unknown to us and exhibits extraordinary high nonspecific staining levels under even rigorous staining protocols due to its unmasked charges (Supplementary Figure S11). Other dyes yielded the desired low nonspecific staining and controlled blinking behavior. Nevertheless, dyes with masked charges (e.g. CF680) never achieved high labeling efficiencies, which may be partially attributed to their increased size (e.g. CF680 has about 2.3 times the molecular weight of Alexa Fluor 680 due to the masking groups) and the high molecular crowding of *S. pombe* cells (Supplementary Figure S11). Among all tested dyes, the best working ones available to us were JF549 and CF647 as seen in Supplementary Figure S11. Nevertheless, for all labeling strategies involving dyes, achieving reliable and high labeling efficiencies as well as quantitative dual-color SMLM read-outs without any channel crosstalk or a high miss rate of events (we tested chromatic dual-color, spectral demixing and dye-FP-mixture strategies) remained problematic. Thus, all measurement strategies in our final selection solely relied on direct FP fusions and were all in the orange-red part of the visual spectrum. As mEos2, our first FP of choice induced conspicuous phenotypical anomalies for several POIs (and is known to artificially aggregate and self-oligomerize at higher densities interfering with POI positioning and function (Wang et al., 2014)), we carefully tested different FPs and ultimately decided to utilize a dual-color approach published in Virant et al. (Virant et al., 2017), which relies on mEos3.2-A69T and PAmcherry1 and added mScarlet-I as the label for the SPB reference.
