## Supplementary stan code and data for distance calculations for "Unraveling the kinetochore nanostructure in *Schizosaccharomyces pombe* using multi-color single-molecule localization microscopy": kinetochore.stan

```
data {

    // number of data points
    int<lower=0> N;
    
    // measured distances, A = cnp1, B = POI, C1 = spindle pole 1, C2 = spindle pole 2
    vector<lower=0>[N] AB;
    vector<lower=0>[N] BC1;
    vector<lower=0>[N] BC2;
    vector<lower=0>[N] AC1;
    vector<lower=0>[N] AC2;
    vector<lower=0>[N] C1C2;

    // estimated variance of Gaussian measurement error of the spindle poles
    vector<lower=0>[N] var_C1;
    vector<lower=0>[N] var_C2;

    // number of movies
    int<lower=0> N_movies;
    
    // source movie for all data points
    int<lower=1,upper=N_movies> movie_id[N];

    // tau square an nu parameters for inverse chi square prior for the alignment error, per movie
    // alignment between channels A (cnp1) and B (POI)
    vector<lower=0>[N_movies] tau_square_alignment_B;
    vector<lower=1>[N_movies] nu_alignment_B;
    // alignment between channels A (cnp1) and C (spindle poles)
    vector<lower=0>[N_movies] tau_square_alignment_C;
    vector<lower=1>[N_movies] nu_alignment_C;

    // tau square an nu parameters for inverse chi square prior for the cluster error
    real<lower=0> tau_square_cluster;
    int<lower=1> nu_cluster;
}


transformed data {

    // sigma for weakly informative Gaussian prior for AB
    real sigma_prior_AB =  mean(AB) * 10;
    
    // mixture coefficient, i.e. prior probability that cnp1 and POI are attached to spindle pole 1
    vector[N] lambda = 1 ./ (1 + (AC2 ./ AC1).^-3);


    vector[N] var_C[2] = {var_C1, var_C2};


    // Construct two triangles with the given distances such that
    // - A lies on the y axis and above the x axis
    // - C1 and C2 lie on the x axis
    
    vector[N] A[2];
    vector[N] B[2];
    vector[N] C[2, 2];

    // start out with C1 on the y axis
    C[1][1] = zeros_vector(N);
    C[1][2] = zeros_vector(N);
    C[2][2] = zeros_vector(N);
    C[2][1] = C[1][1] + C1C2;
    {
        vector[N] phi_A = acos((AC1.^2 + C1C2.^2 - AC2.^2) ./ (2 * AC1 .* C1C2));   // angle C2C1A
        A = {AC1 .* cos(phi_A), AC1 .* sin(phi_A)};
        vector[N] phi_B = acos((BC1.^2 + C1C2.^2 - BC2.^2) ./ (2 * BC1 .* C1C2));   // angle C2C1B
        B = {BC1 .* cos(phi_B), BC1 .* sin(phi_B)};

        // If the distance between A and B is closer to param AB if B is reflected through the x axis, do that.
        // (In theory, the distance should be exactly AB in one of the two cases, but rounding issues.)
        vector[N] d1 = fabs(sqrt((A[1]-B[1]).^2 + (A[2]-B[2]).^2) - AB);
        vector[N] d2 = fabs(sqrt((A[1]-B[1]).^2 + (A[2]+B[2]).^2) - AB);
        for(i in 1:N) {
            if(d1[i] > d2[i]) {
                B[2][i] *= -1;
            }
        }
    }

    // move A onto the y axis
    B[1]    -= A[1];
    C[1][1] -= A[1];
    C[2][1] -= A[1];
    A[1]    -= A[1]; 
}


parameters {

    // "raw", centered parameters, need to be translated and scaled to yield actual values (see below und "transformed paramters")

    real AB_true_raw;

    vector[N] A_true_raw[2, 2];
    vector[N] C_true_raw[2, 2];

    vector<lower=0>[N] var_A_raw;
    vector<lower=0>[N] var_B_raw;
    vector<lower=0>[N_movies] var_alignment_B_raw;
    vector<lower=0>[N_movies] var_alignment_C_raw;
    
    
    // sigma for hierarchical, uninformative Gaussian prior for positions of A, C1 and C2
    real<lower=0> sigma_prior;
}


transformed parameters {

    // translate and scale "raw" parameter values to yield actual values

    real AB_true = AB_true_raw * sigma_prior_AB;
    
    vector[N] A_true[2, 2];
    vector[N] B_true[2, 2];
    vector[N] C_true[2, 2];
    for(i in 1:2) {
        A_true[i][1] = A_true_raw[i][1] * sigma_prior + A[1];
        A_true[i][2] = A_true_raw[i][2] * sigma_prior + A[2];
        C_true[i][1] = C_true_raw[i][1] * sigma_prior + C[i][1];
        C_true[i][2] = C_true_raw[i][2] * sigma_prior + C[i][2];

        vector[N] v[2] = {C_true[i][1]-A_true[i][1], C_true[i][2]-A_true[i][2]};
        vector[N] fac = AB_true ./ sqrt(v[1].^2 + v[2].^2);
        B_true[i][1] = A_true[i][1] + v[1] .* fac;
        B_true[i][2] = A_true[i][2] + v[2] .* fac;
    }
    
    
    // combine measurement, alignment and cluster errors to calculate individual errors for all positions

    vector[N] sigma_A;
    vector[N] sigma_B;
    vector[N] sigma_C[2];
    {
        vector[N] var_A = var_A_raw * nu_cluster * tau_square_cluster;
        vector[N] var_B = var_B_raw * nu_cluster * tau_square_cluster;
        vector[N_movies] var_alignment_B = var_alignment_B_raw .* nu_alignment_B .* tau_square_alignment_B;
        vector[N_movies] var_alignment_C = var_alignment_C_raw .* nu_alignment_C .* tau_square_alignment_C;
        
        sigma_A = sqrt(var_A);
        for(i in 1:N) {
            sigma_B[i]    = sqrt(var_B[i]    + var_alignment_B[movie_id[i]]);
            sigma_C[1][i] = sqrt(var_C[1][i] + var_alignment_C[movie_id[i]]);
            sigma_C[2][i] = sqrt(var_C[2][i] + var_alignment_C[movie_id[i]]);
        }
    }
}


model {

    // priors; as the parameters are centralized, we use standard distributions here

    AB_true_raw ~ std_normal();
    
    for(i in 1:2) {
        for(j in 1:2) {
            A_true_raw[i][j] ~ std_normal();
            C_true_raw[i][j] ~ std_normal();
        }
    }

    var_A_raw ~ inv_chi_square(nu_cluster);
    var_B_raw ~ inv_chi_square(nu_cluster);
    var_alignment_B_raw ~ inv_chi_square(nu_alignment_B);
    var_alignment_C_raw ~ inv_chi_square(nu_alignment_C);
    
    
    // for both spindle poles, calculate probability of measuring A, B, and C if true positions are A_true, B_true and C_true,
    // assuming Gaussian measurement errors with sigmas sigma_A, sigma_B and sigma_C
    // then calculate combined probability for both options assuming a prior probability of lambda for option 1

    for(i in 1:N) {    
        real lp[2] = {0.0, 0.0};
        for(j in 1:2) {
            for(k in 1:2) {
                lp[j] += normal_lpdf(A[k][i]    | A_true[j][k][i], sigma_A[i]);
                lp[j] += normal_lpdf(B[k][i]    | B_true[j][k][i], sigma_B[i]);
                lp[j] += normal_lpdf(C[j][k][i] | C_true[j][k][i], sigma_C[j][i]); 
            }
        }
        target += log_mix(lambda[i], lp[1], lp[2]);
    }
}
```
