## Supplementary material for "Unraveling the kinetochore nanostructure in *Schizosaccharomyces pombe* using multi-color single-molecule localization microscopy": Example csv SMLM localization data: info.txt

We showcase two examples for our annotation using two csvs of annotated SMLM localization data of cells with a mitotic spindle that passed the quality checks of drift control and channel overlay. Importantly, all data analysis steps were conducted on the localization data directly (see data analysis), images were only used for image representation purposes.
For the first example, a cell with dam1 as POI, all 6 kinetochores are visible and all were grouped and separated from noise. One can also nicely see (due to the large distance of dam1 to cnp1) which of the six belong to which spindle by the spatial orientation of the cnp1 and dam1 cluster to each other, and both groups of three have a pair that is spatially closer to the wrong pole, important for the distance calculations. The second example, with spc7 as the example POI, has only 5 visible clusters. A closer look shows that one of them has unusual dimensions and a high number of localizations, representing most likely two overlapping centromeric regions. Thus, for this cell, only 4 kinetochore pairs were annotated for further analysis. Also, the cluster pairs are overlapping much more, so a direct decision to which spindle they belong to gets difficult. 
