## Supplementary figures and images for "Unraveling the kinetochore nanostructure in *Schizosaccharomyces pombe* using multi-color single-molecule localization microscopy"

### S1.png

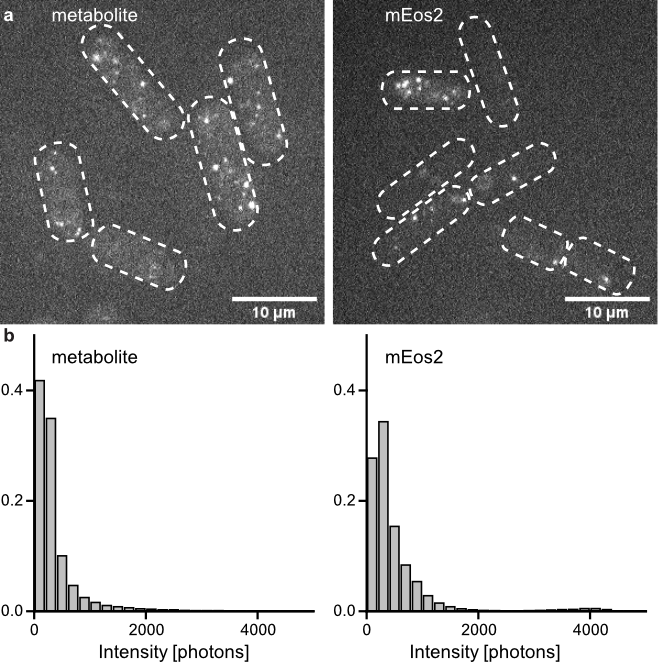

### S2.png

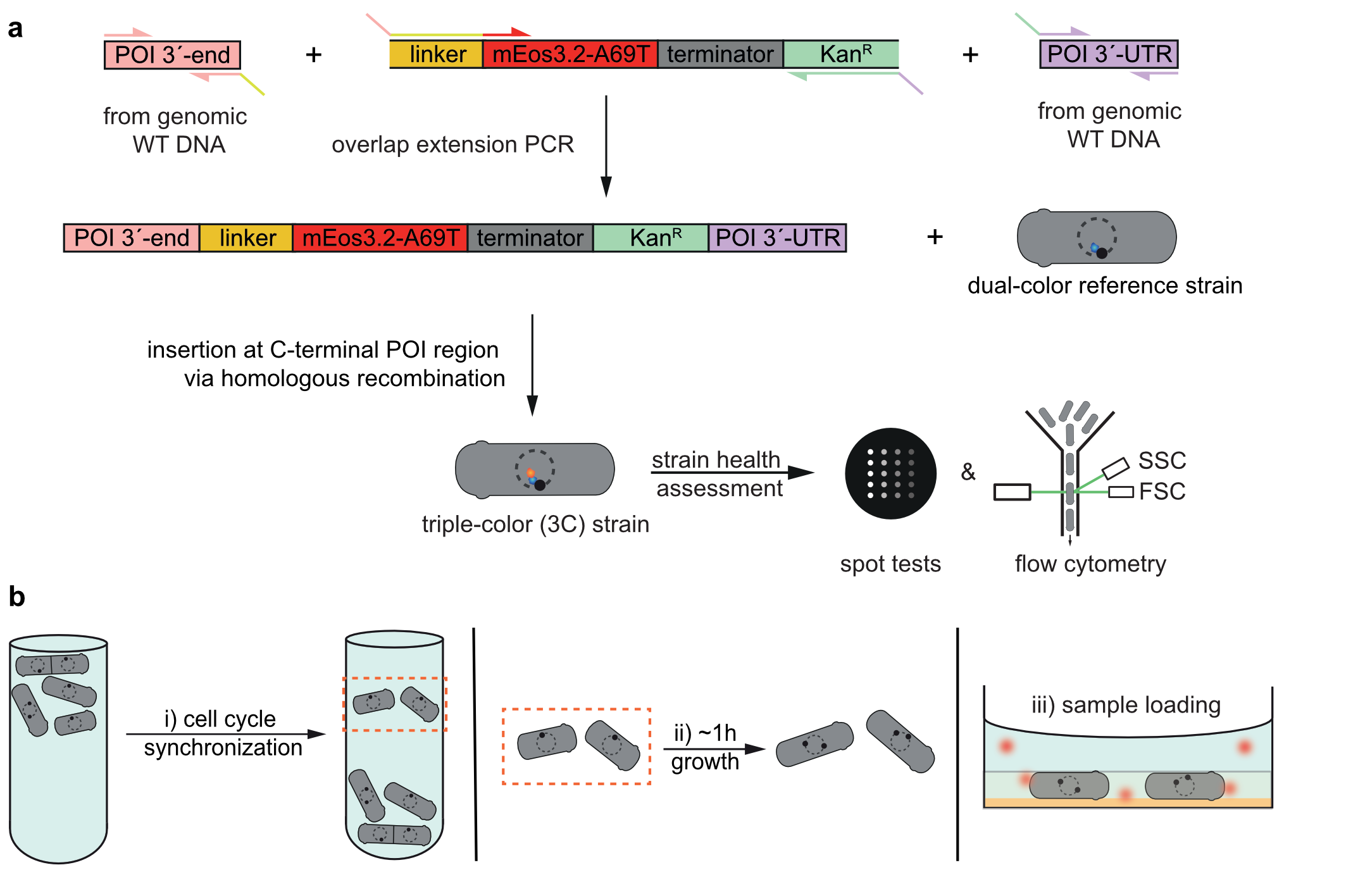

### S3.png

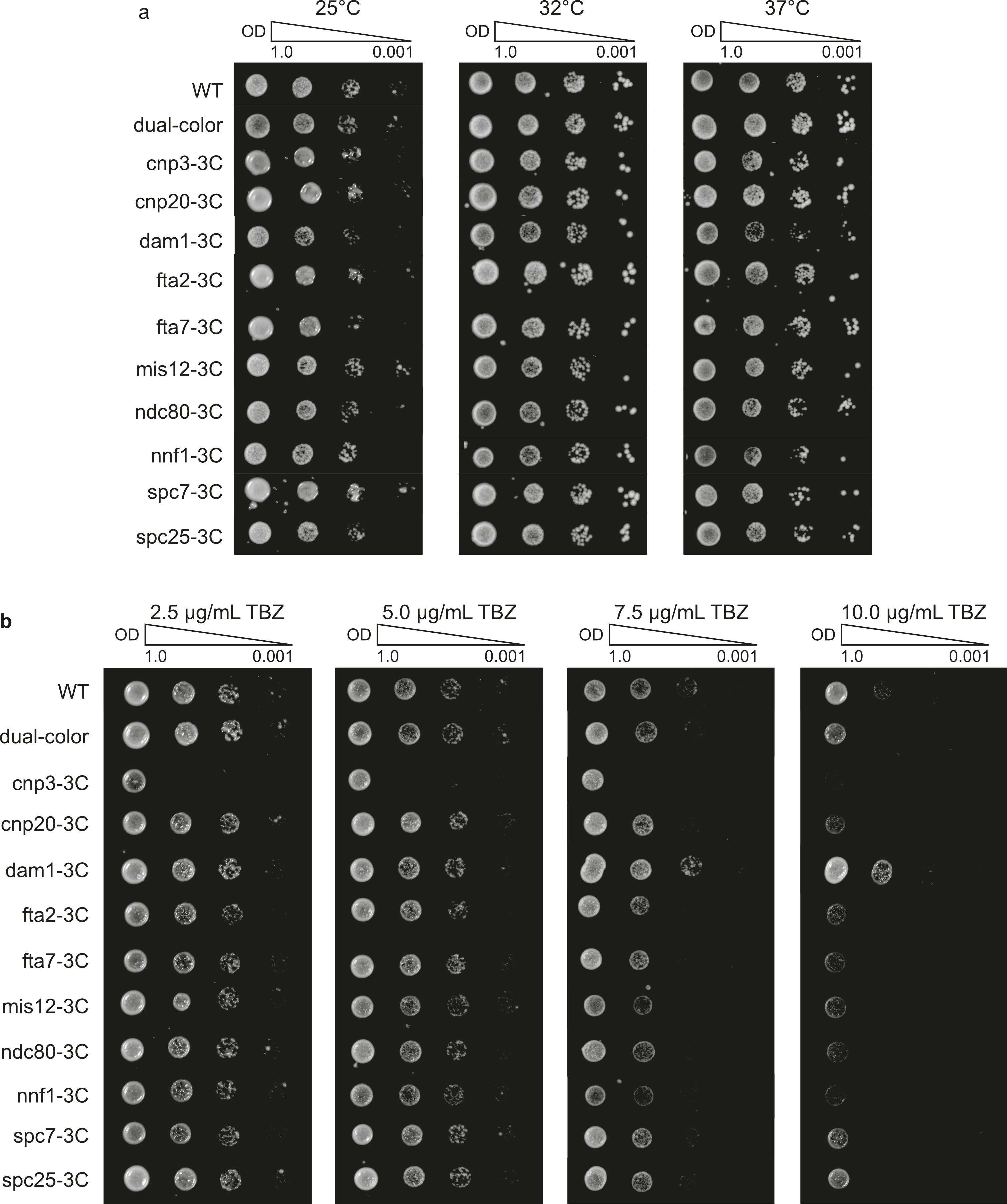

### S4.png

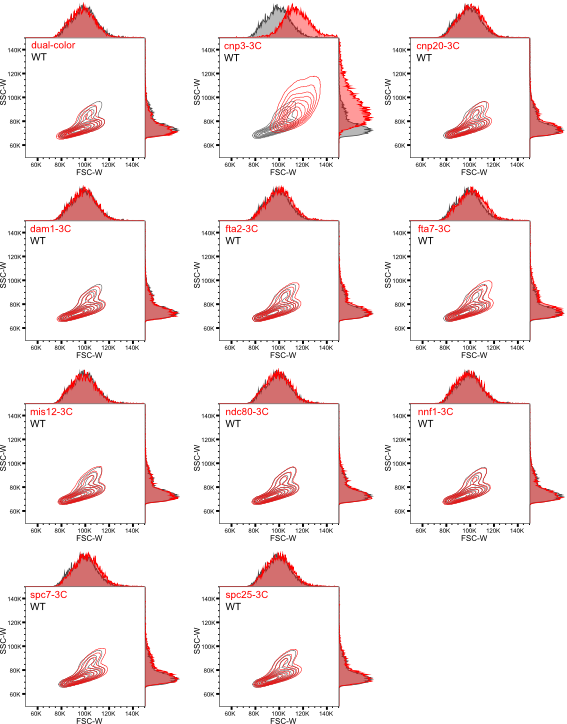

### S5.png

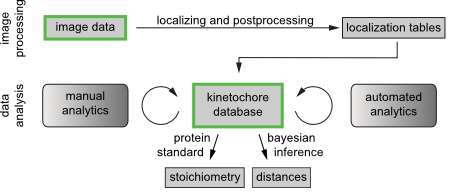

### S6.png

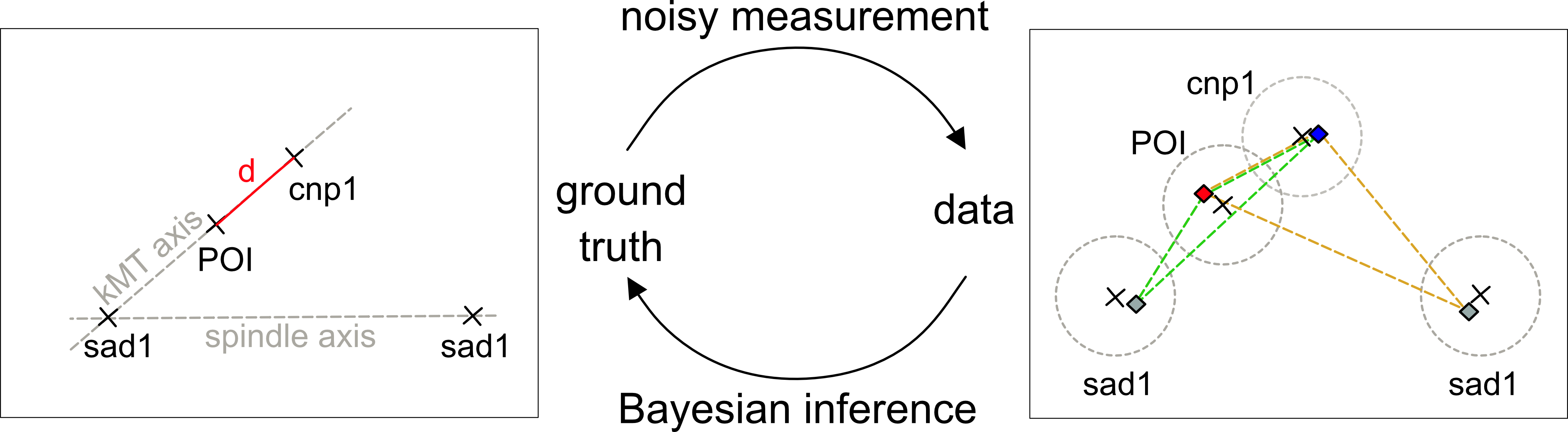

### S7.png

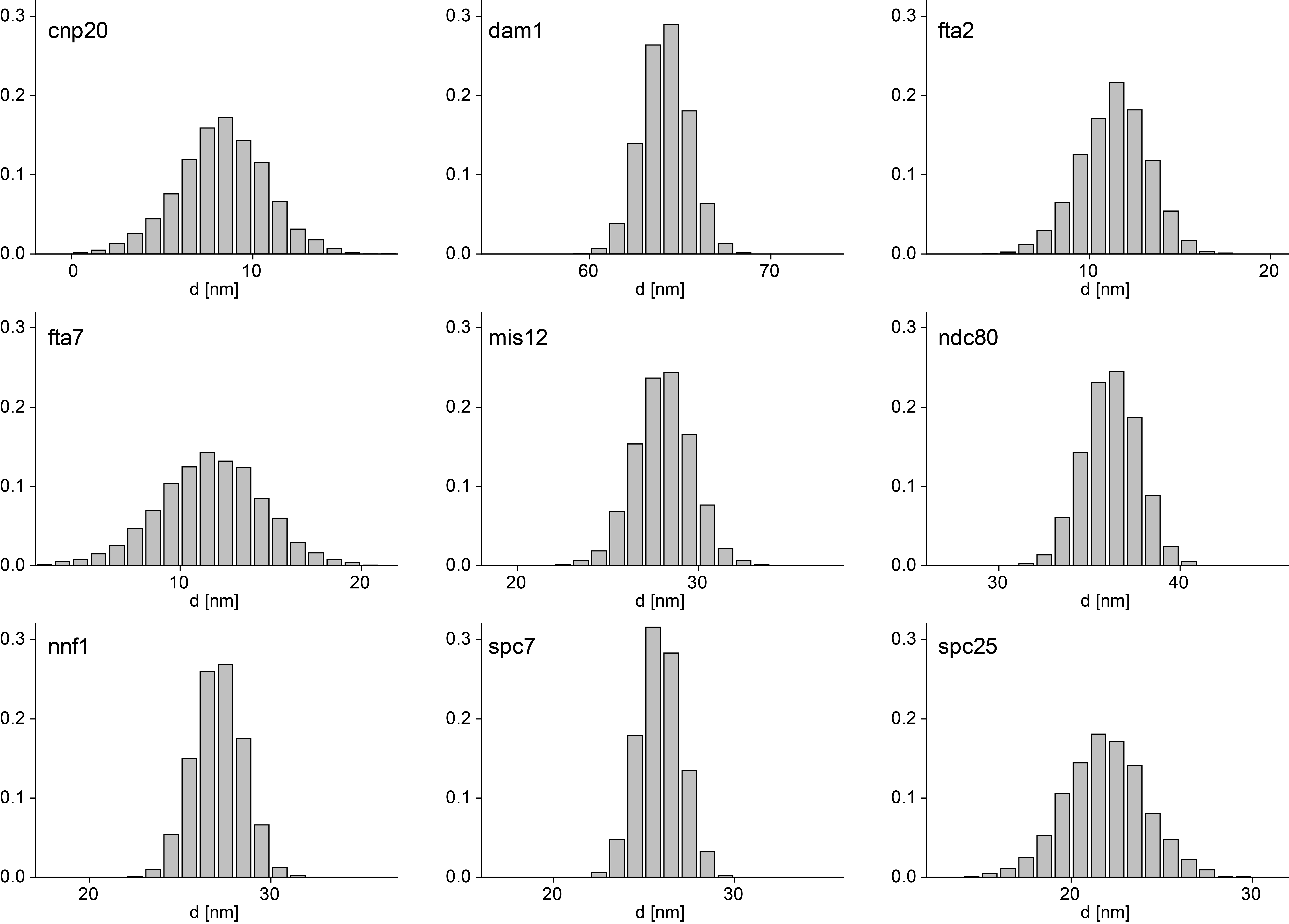

### S8.png

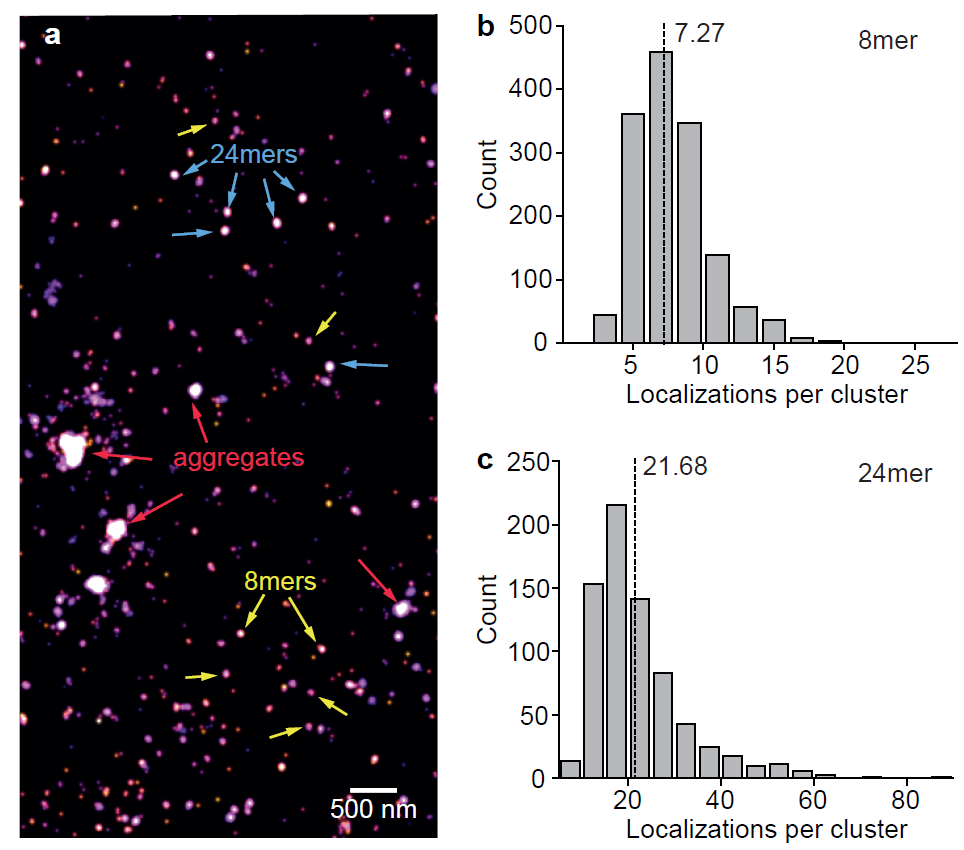

### S9.png

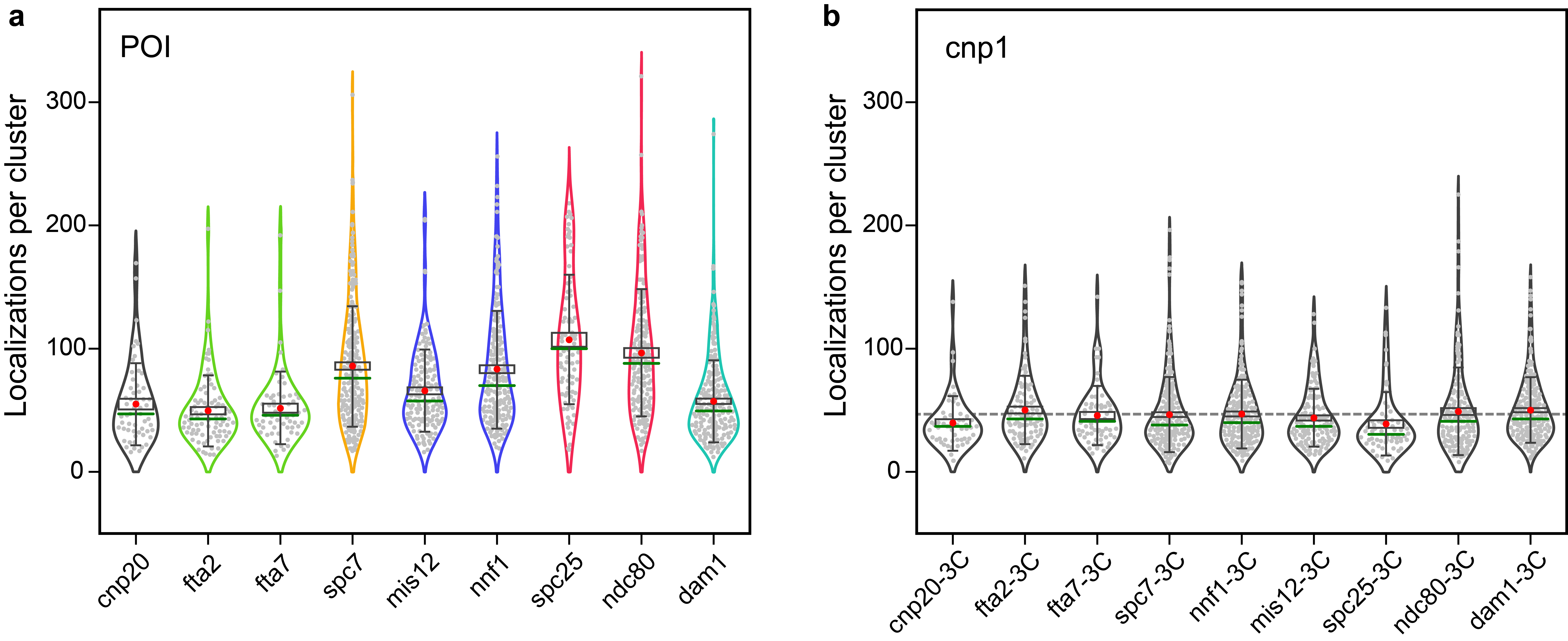

### S10.png

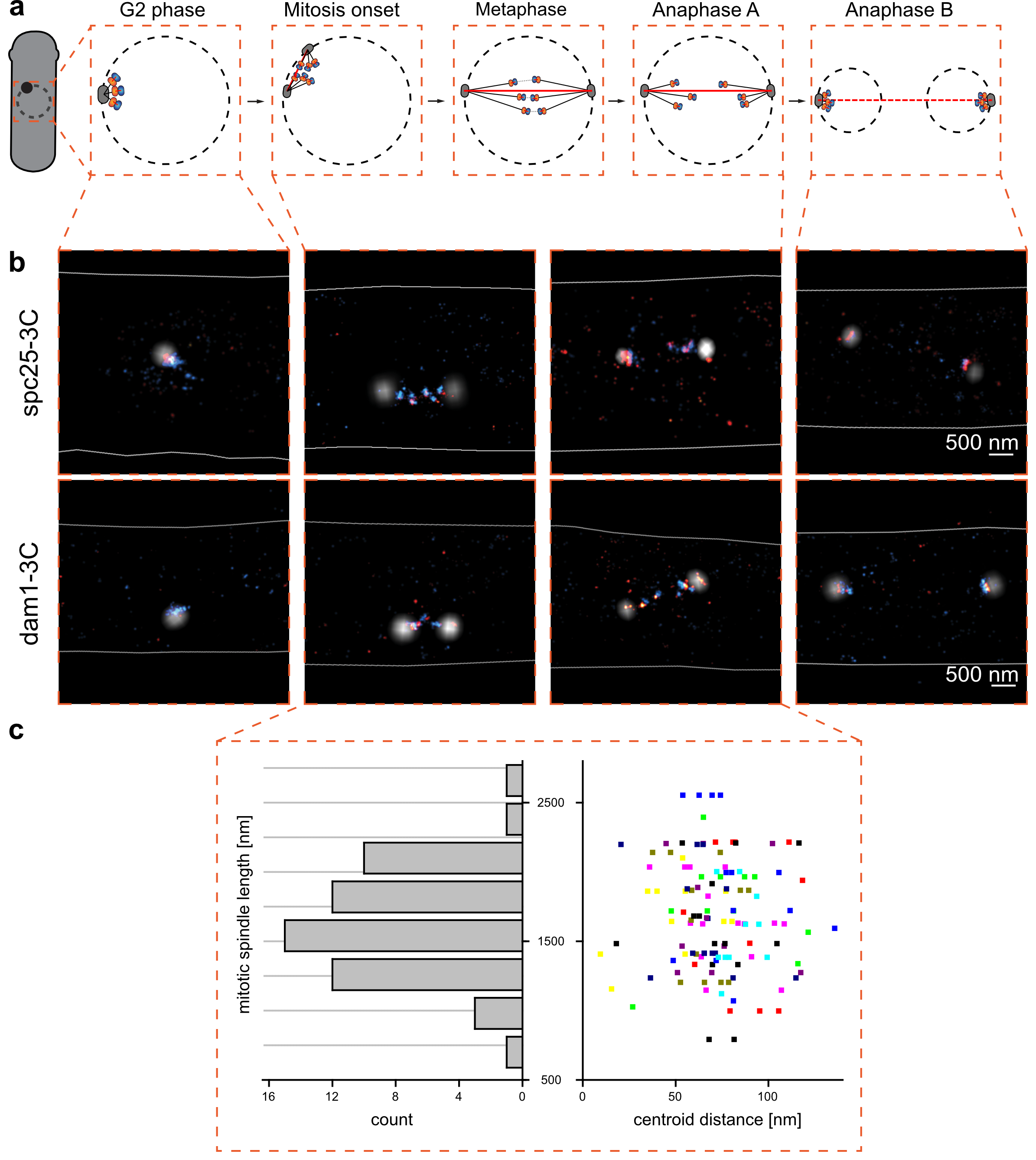

### S11.png

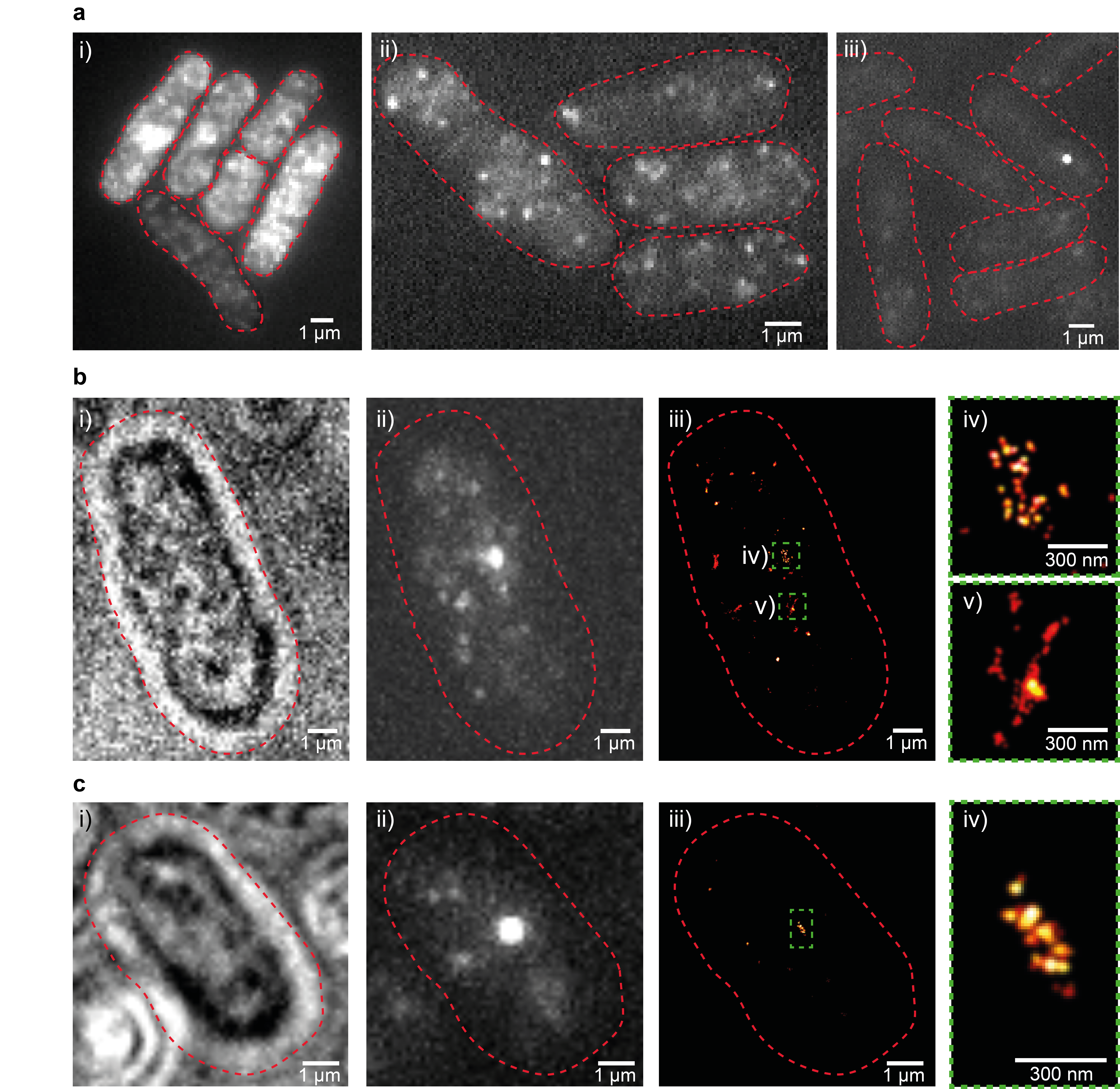

### S12.png

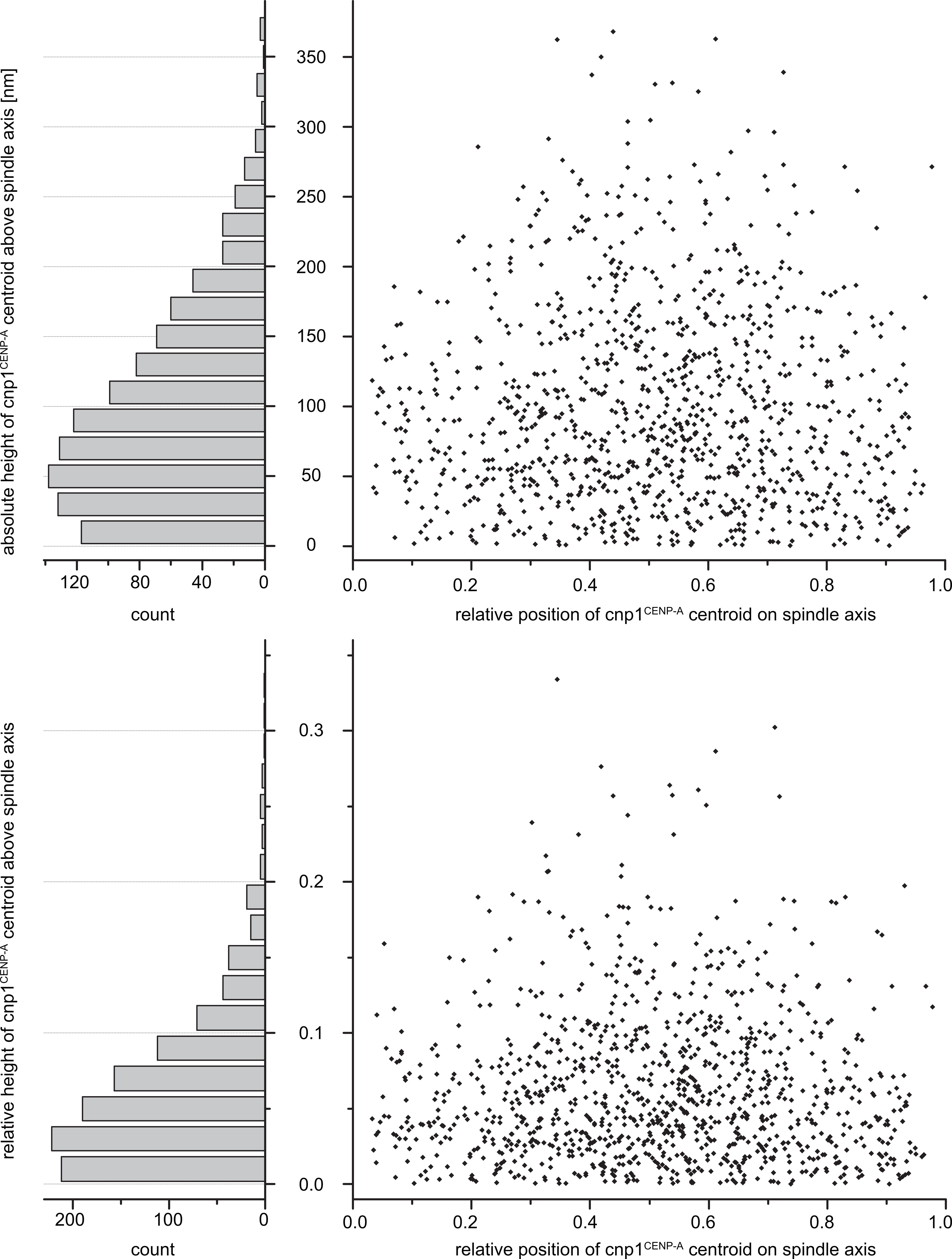
